# Continuous negative autoregulation fine-tunes dosage-sensitive transcription factor expression to maintain post-mitotic neuron identity

**DOI:** 10.64898/2026.05.14.725191

**Authors:** Honorine Destain, Alan Koh, Heewhan Shin, André E. X. Brown, Paschalis Kratsios

## Abstract

How post-mitotic neurons maintain precise transcription factor (TF) levels throughout life remains a fundamental open question. Here, we challenge the prevailing model of positive autoregulation by demonstrating that UNC-3 (Collier/EBF1-4), a dosage-sensitive TF continuously required for cholinergic motor neuron identity in *C. elegans*, negatively regulates its own expression. Using genetics, biochemistry, and inducible protein depletion, we show this self-repression occurs directly at the transcriptional level and persists beyond development. CRISPR/Cas9 disruption of negative autoregulation causes motor neuron identity and locomotion defects, establishing its functional necessity. Mechanistically, the UNC-3 DNA-binding domain is required and sufficient for self-repression, with an AlphaFold2 screen implicating chromatin factors as interaction partners. Critically, UNC-3 self-repression is continuously counterbalanced by positive input from the HOX cofactor CEH-20/PBX, revealing a dynamic “balancing act” between opposing regulatory inputs that stabilize TF dosage over time. Mutations in the *unc-3* ortholog *EBF3* cause a neurodevelopmental syndrome, and disease-associated variants disrupt UNC-3 self-repression, revealing a key molecular mechanism underlying the disorder. We propose that negative autoregulation continuously counteracted by positive input represents a broadly applicable principle for maintaining dosage-sensitive TF expression to secure post-mitotic cell identity.

## INTRODUCTION

Cell type identity and thus organismal phenotypes are often sensitive to transcription factor (TF) levels ^1–5^. Changes in TF levels, caused either by copy number variation or mutations in *cis*-regulatory sequences, are implicated in human disease ^6^. For example, both increased and decreased copy number of *EBF3* have been linked to a neurodevelopmental syndrome termed HADDS (Hypotonia, Ataxia, and Delayed Development Syndrome) ^7,8^. Additionally, *cis-*regulatory mutations in the homeodomain TF NKX2-5 can lead to either increased or decreased NKX2-5 levels that impact heart development ^9,10^. Overall, a large body of work indicates that TF levels must be precisely regulated during animal development to ensure the appropriate generation of cell types and tissues.

In development, appropriate induction of TF expression in specific cell types is often regulated by signaling molecules and transiently expressed TFs ^11,12^. Following their initial launching in post-mitotic cell types, certain TFs maintain their expression for extended periods of time to ensure a particular cell fate and function ^13–15^. The importance of maintenance of TF levels is highlighted by accumulating evidence suggesting that failure to maintain TF levels can cause disease. For example, decreased expression of *NURR1* in midbrain dopamine neurons has been observed in Parkinson’s disease, an adult-onset neurological disorder ^16^.

TFs function within gene regulatory networks (GRNs), which represent the sum of TF and target gene interactions, mediated via *cis*-regulatory regions ^17^. In development, the outcome of GRNs is the expression of a unique set of effector genes, which endow individual cell types with their unique molecular identities and specialized functions. GRNs are composed of network motifs – statistically enriched TF interactions serving specific biological functions ^18^. Decades of developmental genetics in invertebrate and vertebrate models have elucidated TF network motifs in early embryonic development ^17,19^. However, such studies mostly utilized constitutive approaches, preventing temporal dissection of TF function. As a consequence, the network motifs that function over time to maintain TF expression in adult post-mitotic cell types are poorly understood.

Which network motifs maintain TF expression in adult cell types? The simplest motif is autoregulation – regulation of a TF’s gene expression by its own protein product. Autoregulation can be either positive or negative ^20^. Positive autoregulation can function as a ‘lock-in’ device to maintain TF expression and cell identity even after the initial activating signal has petered out ^21^. The role of positive autoregulation has been extensively demonstrated - both computationally and experimentally across many biological contexts ^20,22–27^. Disruption of positive autoregulation can cause disease; for example, mutation of *cis*-regulatory elements essential for positive autoregulation of *PAX6* disrupts maintenance of its expression and causes aniridia, a condition characterized by the partial or complete absence of the iris ^28^. On the other hand, negative TF autoregulation can reduce cell-to-cell variability in TF expression levels, but this function has primarily been studied in prokaryotes (λ phage, *E. coli*) ^29,30^. Disruption of negative TF autoregulation also causes disease, exemplified by *cis*-regulatory mutations in the proto-oncogene *BCL6* that cause lymphoma ^31,32^. Compared to positive autoregulation though, the mechanisms underlying negative TF autoregulation remain largely unknown, especially in the context of long-lived postmitotic cells.

Neurons are postmitotic cells that must maintain their identity and function throughout life, offering the opportunity to identify GRNs that control the identity of long-lived cell types. Neuronal cell type identity is established during development, and maintained in adulthood, by a class of TFs known as terminal selectors ^13^. Terminal selectors are continuously expressed in specific neuron types, from development to adulthood, where they directly activate effector genes necessary for neuronal identity and function ^13^. To date, over a hundred terminal selectors have been described for specific neuron types in nematodes (*C. elegans*) ^33^, fruit flies (*Drosophila)* ^34^, planarians (*Schmidtea mediterranea*) ^35^, cnidarians (*Nematostella vectensis)* ^36^, marine chordates (*Ciona robusta)* ^37^, zebrafish (*Danio rerio*) ^38^, and mice (*Mus musculus*) ^39^, indicating a deeply conserved role for these critical regulators of neuronal identity. They are biomedically relevant; mutations in terminal selector orthologues are linked to both neurodevelopmental and neurodegenerative conditions ^40,41^.

Studies of a handful of terminal selectors in nematodes and mice (*ttx-3, ceh-10, che-1, Pet-1*) demonstrated that their expression is induced during development by transient inputs from TFs and classic signaling pathways, such as Wnt ^42–44^. After initial induction, terminal selector expression is thought to be maintained via positive autoregulation, based on studies conducted on nine terminal selectors to date (*mec-3, che-1, odr-7, ttx-3, cfi-1, lin-39, Pet-1, Glass,* and *hlh-15*) ^22,34,39,42,45–54^. Although positive autoregulation has emerged as an important molecular principle, additional mechanisms are required to stably maintain terminal selector expression and thereby cell fate ^50,55–57^; these mechanisms remain poorly understood.

Here, we study the gene regulatory mechanisms that fine-tune the expression of a dosage-sensitive and clinically relevant TF in post-mitotic neurons. We use as a paradigm UNC-3 (Collier/EBF1-4), a terminal selector of cholinergic motor neuron (MN) identity in the *C. elegans* ventral nerve cord, where it is known to act as transcriptional activator^58–63^. We found that UNC-3 directly and continuously negatively autoregulates to repress its own expression levels in MNs, challenging the current model of positive autoregulation for terminal selector-type TFs. Our results are clinically relevant because altered expression of the *unc-3* human ortholog, *EBF3*, causes hypotonia, ataxia, and delayed development syndrome (HADDS), a neurodevelopmental syndrome with motor symptoms^7,8,64^. We found that HADDS-causing *EBF3* variants disrupt *unc-3* negative autoregulation when tested in *C. elegans*. Mechanistically, we show that the DNA binding and Immunoglobulin/Plexin/TF (IPT) domains of UNC-3 are required for self-repression. With an *in silico* screen, we identify TFs and chromatin-associated factors predicted to interact with UNC-3 to modulate its function. Importantly, negative autoregulation is continuously counterbalanced by opposing activating input from the conserved HOX cofactor CEH-20/PBX. Altogether, we propose that regulation of dosage-sensitive TFs over time through a ‘balancing-act’ of continuous self-repressing and activating inputs may constitute a general principle for cell fate maintenance in long-lived cell types.

## RESULTS

### *unc-3* negatively autoregulates in cholinergic nerve cord motor neurons

Given the nine documented cases of positive autoregulation of terminal selectors to date ^22,34,39,42,45–54^, we first sought to determine whether *unc-3* has any autoregulatory input. To assess this, we generated a ∼30kb transgenic reporter for *unc-3* (*unc-3_Exon2_mCherry*) that contains all the *cis*-regulatory information upstream and downstream of the *unc-3* locus, but the region spanning exons 3 to 12 has been replaced with *mCherry* (**Fig. 1A**) (see Methods). Next, we crossed this reporter into two *unc-3* loss-of-function (LoF) mutants (W309*, R166W); W309* is a presumptive null that introduces a premature stop codon in IPT domain^65^, and R166W is a missense mutation in the Zinc finger ^65,66^ (**Fig. 1A**). Notably, R166W is one of the *de novo* mutations that cause the *EBF3* neurodevelopmental syndrome (HADDS) (**Fig 4A**) ^67^. Compared to control animals, we observed increased *unc-3_Exon2_mCherry* fluorescence intensity in cholinergic MNs of both mutants at the fourth larval stage (L4), after both waves of MN generation are complete (**Fig. 1B, D-E**). This effect was specific to cholinergic MNs (**Supp. Fig. 1A**). Increased *unc-3* reporter expression was also observed at the first larval stage (L1), after the first wave of MN generation is complete (**Fig. 1B-C, E**). These findings suggest that, in contrast to other terminal selectors, *unc-3* negatively, rather than positively, autoregulates, and this negative autoregulation is established early in development, at least by L1.

**Figure 1:**
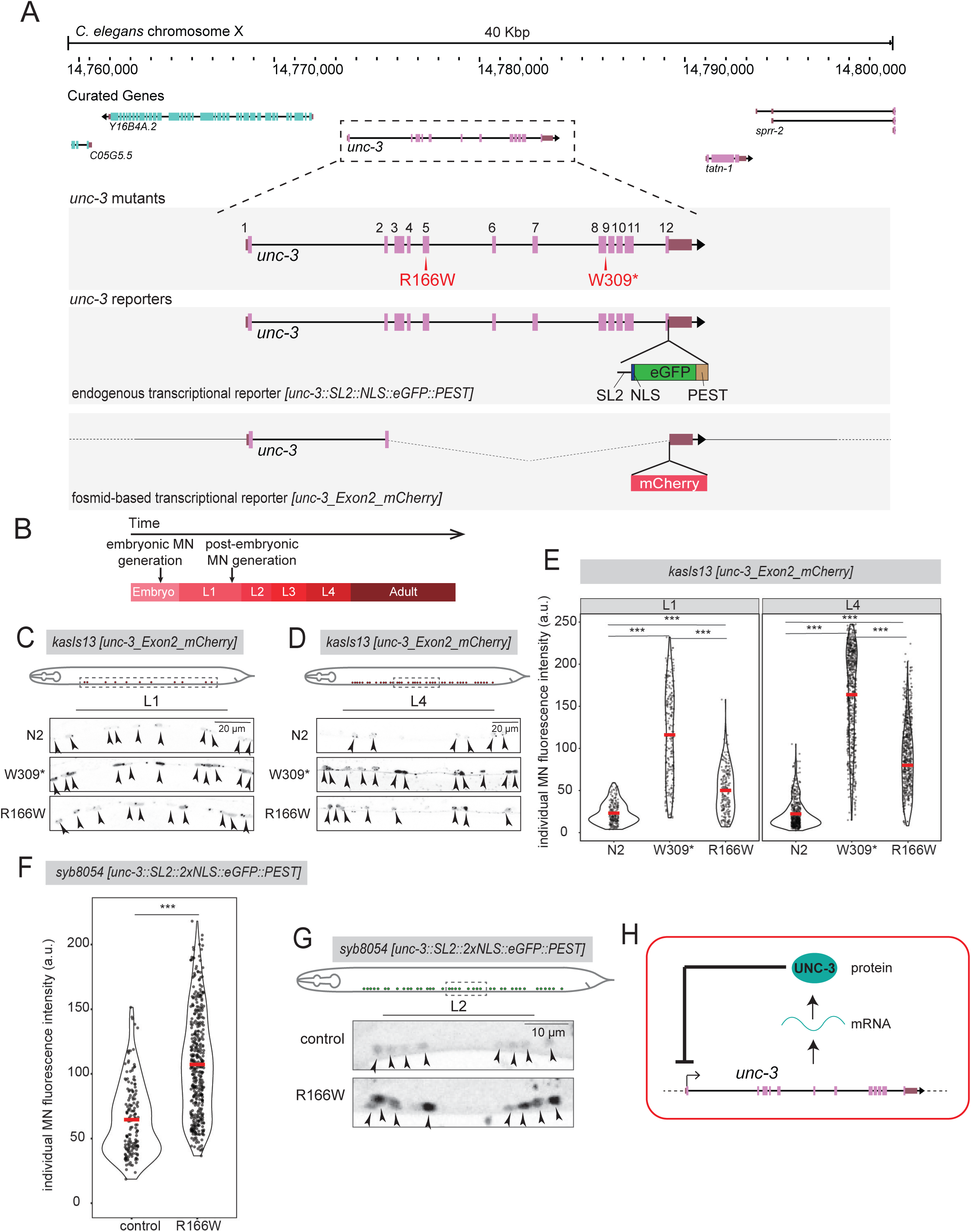
*unc-3* negatively autoregulates in *C. elegans* motor neurons. **A:** Genomic view of the endogenous *unc-3* locus, with zoomed in views depicting (top) *unc-3* mutants (in red), and (bottom) *unc-3* endogenous transcriptional reporter *[unc-3::SL2::NLS::eGFP::PEST]* and transgenic fosmid-based transcriptional *unc-3* reporter *[unc-3_Exon2_mCherry]* **B:** Schematic of timing of motor neuron (MN) generation in *C. elegans* development. **C:** Representative images of fosmid-based transcriptional *unc-3* reporter (*kasIs13 [unc-3_Exon2_mCherry]*) expression at L1 stage, in N2, *unc-3(W309*)*, and *unc-3(R166W)* backgrounds. Top: Schematic of nerve cord section displayed in images. Arrowheads: motor neurons. Scale bar: 20 µm **D:** Representative images of fosmid-based transcriptional *unc-3* reporter (*kasIs13 [unc-3_Exon2_mCherry]*) expression at L4 stages, in N2, *unc-3(W309*)*, and *unc-3(R166W)* backgrounds. Top: Schematic of nerve cord section displayed in images. Arrowheads: motor neurons. Scale bar: 20 µm **E:** Quantification of individual motor neuron fluorescence intensity at L1 (left) and L4 (right) of transgenic fosmid-based transcriptional *unc-3* reporter (*kasIs13 [unc-3_Exon2_mCherry]*) in N2, *unc-3(W309*)*, and *unc-3(R166W)* backgrounds. Dots represent individual motor neurons. Red bar represents mean. Dunn Kruskal-Wallis multiple comparisons test with Bonferroni correction. (L1: N2 n = 22 worms, W309* n = 20, R166W n = 20; L4 N2 n = 20, W309* n = 20, R166W = 18) **F:** Quantification of individual motor neuron fluorescence intensity at L2 of *unc-3* endogenous transcriptional reporter *[unc-3::SL2::NLS::eGFP::PEST]* in control and *unc-3(R166W)* backgrounds. Dots represent individual motor neurons. Red bar represents mean. Welch’s T-test. (control n = 6 worms, *unc-3(R166W)* n = 12) **G:** Representative images of *unc-3* endogenous transcriptional *[unc-3::SL2::NLS::eGFP::PEST]* reporter at L2 stages in N2 and R166W backgrounds. Top: Schematic of nerve cord section displayed in images. Arrowheads: motor neurons. Scale bar: 10 µm **H:** Model of transcriptional negative autoregulation occurring at the *unc-3* endogenous locus. For all quantifications, p-values: * < .05, ** <.01, *** < .001, n.s. = no significance.

To exclude the possibility that negative autoregulation is observed due to the transgenic nature of the *unc-3_Exon2_mCherry* reporter, we next generated an endogenous *eGFP* transcriptional reporter, *unc-3::SL2::NLS::eGFP::PEST*. The bicistronic SL2 linker sequence leads to separation of the *unc-3* and *NLS::eGFP::PEST* mRNA sequences ^68^, allowing eGFP fluorescent intensity to reflect *unc-3* transcript levels (**Fig. 1A**). Destabilization by the PEST tag significantly reduces eGFP protein half-life, increasing the temporal resolution of transcriptional dynamics by preventing eGFP perdurance ^69^. Next, we introduced the syndromic R166W Zinc-finger mutation in the context of the endogenous *unc-3::SL2::NLS::eGFP::PEST* reporter (**Fig. 1A**). We observed a significant increase of eGFP expression in cholinergic MNs at L2, demonstrating that *unc-3* negative autoregulation does occur at the endogenous *unc-3* locus, requires an intact Zinc-finger domain, and is affected by a syndromic mutation (**Fig. 1F-G**). Increased expression was also observed at L4 (**Supp. Fig. 1B-C**). Altogether, these data offer the first example of a terminal selector gene that negatively regulates its own expression (**Fig. 1H**).

### *unc-3* negative autoregulation is maintained

The analysis of L1 animals with constitutive *unc-3* LoF alleles (W309*, R166W) indicates that *unc-3’s* negative autoregulation is established early during development (**Fig. 1C-E**). To test whether it continues to occur during post-embryonic and/or adult life stages, we conditionally depleted the UNC-3 protein starting at the L3 stage (∼15 hours after the generation of all cholinergic nerve cord MNs) (**Fig. 2C**). Using the auxin-inducible degron (AID) system ^70^ and an available *unc-3::mNG::3xFLAG::AID* allele ^71^, we induced UNC-3 depletion with the plant hormone auxin in the presence of the plant F-box protein, TIR-1 (**Fig. 2A-B**). Continuous auxin exposure from late L3 stage until the first day (day 1) of adulthood led to robust depletion of UNC-3 (**Fig. 2C-E**). At day 1, we observed increased expression of the *unc-3* transcriptional reporter in cholinergic MNs, demonstrating that UNC-3 represses its own expression after L3 (**Fig. 2F-G**). Together, our L1 and L4 stage analysis with *unc-3* constitutive alleles and our AID analysis suggest that *unc-3* negative autoregulation in cholinergic MNs occurs across multiple time points, both in early and late larval/young adult stages.

**Figure 2:**
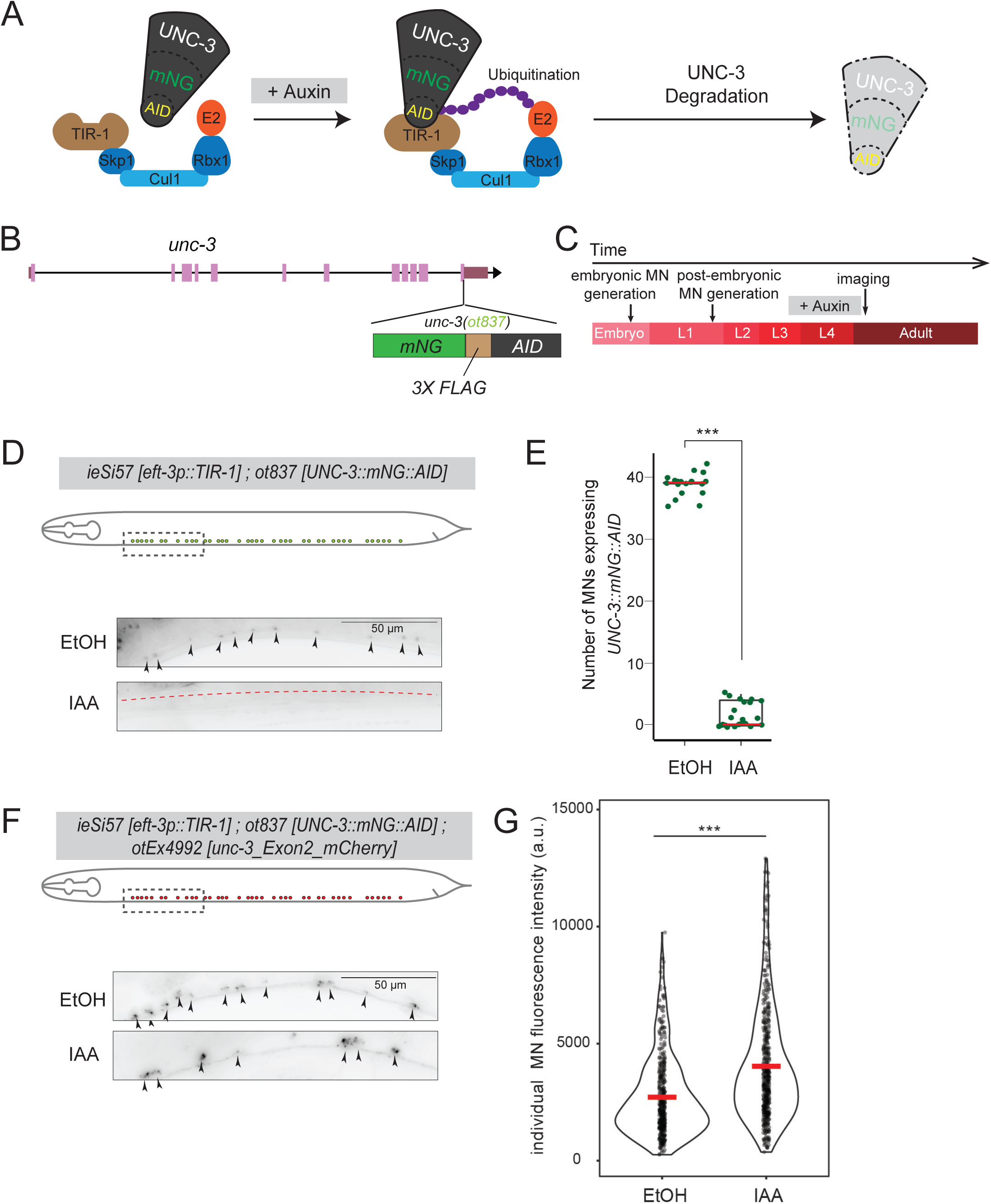
*unc-3* negative autoregulation is continuous. **A:** Schematic of temporally controlled protein depletion with auxin. UNC-3::mNG::AID is recognized by TIR-1 in the presence of auxin, leading to ubiquitination and degradation. **B:** Schematic of endogenously tagged *unc-3* allele with mNeonGreen (mNG), 3X FLAG tag, and auxin-inducible degron (AID). **C:** Schematic of timing of auxin exposure and imaging **D:** Representative images of *unc-3::mNG::AID* expression in day 1 adults upon exposure to control (EtOH) or auxin. Top: Schematic of nerve cord section displayed in images. Arrowheads: motor neurons. Red dashed line delineates gut autofluorescence. Scale bar: 50 µm **E:** Quantification of number of mNG positive nerve cord motor neurons in day 1 adults upon exposure to control (EtOH) or auxin. Dots represent individual worms. Red bar represents median. Welch’s T-test. (EtOH n = 21 worms, auxin n = 23) **F:** Representative images of fosmid-based transcriptional *unc-3* reporter (*[otEx4992 [unc-3_exon2_mCherry]*) in day 1 adults upon exposure to control (EtOH) or auxin. Top: Schematic of nerve cord section displayed in images. Arrowheads: motor neurons. Scale bar: 50 µm **G:** Quantification of individual motor neuron fluorescence intensity of fosmid-based transcriptional *unc-3* reporter in day 1 adults upon exposure to control (EtOH) or auxin. Dots represent individual motor neurons. Red bar represents mean. Welch’s T-test (EtOH n = 19 worms, auxin n = 18) For all quantifications, p-values: * < .05, ** <.01, *** < .001, n.s. = no significance.

### *unc-3* negative autoregulation is direct and involves UNC-3/EBF cognate binding sites

Our analysis of available UNC-3 ChIP-seq data from L2 animals ^61^ demonstrated extensive UNC-3 binding to its own locus (**Fig. 3A**). This binding partially coincides with accessible chromatin based on ATAC-seq peaks from available panneuronal datasets of late L4 animals ^72^ (**Fig. 3A**). Hence, biochemical and chromatin accessibility data support a model where negative autoregulation occurs directly via UNC-3 binding at its own locus.

**Figure 3:**
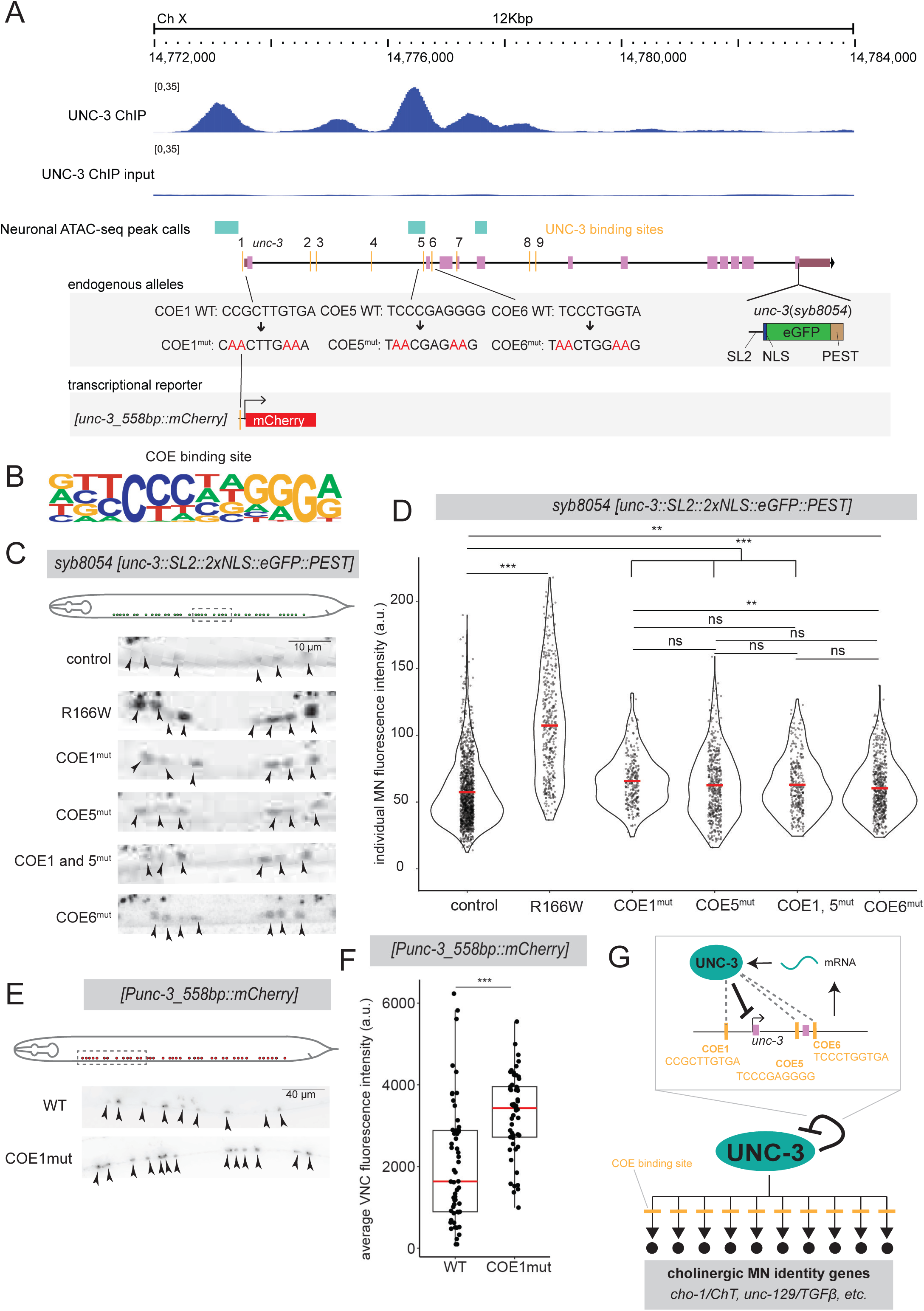
*unc-3* negative autoregulation is direct. **A:** Genomic view of the endogenous *unc-3* locus with UNC-3 ChIP-seq tracks (whole animal, L2, control and input) ^61^ and neuronal ATAC-seq peaks ^72^. Below *unc-3* locus: molecular nature of COE binding site mutations in context of endogenous transcriptional reporter, and transgenic transcriptional reporter. **B:** Positive weight matrix for COE binding site derived from ChIP-seq data using Homer **C:** Representative images of *unc-3* endogenous transcriptional reporter *[unc-3::SL2::NLS::eGFP::PEST]* at L2 in control, *unc-3(R166W),* COE1 mutant, COE5 mutant, COE1 and 5 mutant, and COE6 mutant backgrounds. Top: Schematic of nerve cord section displayed in images. Arrowheads: motor neurons. Scale bar: 10 µm **D:** Quantification of individual motor neuron fluorescence intensity of *unc-3* endogenous transcriptional reporter *[unc-3::SL2::NLS::eGFP::PEST]* at L2 in control, *unc-3(R166W),* COE1 mutant, COE1 deletion, COE5 mutant, COE1 and 5 mutant, and COE6 mutant backgrounds. Dots represent individual motor neurons. Red bar represents mean. Dunn Kruskal-Wallis multiple comparisons test with Bonferroni correction. (control n = 50 worms, *unc-3(R166W)* n = 12, COE1 mutant n = 9, COE5 mutant n = 17, COE1 and 5 mutant n = 14, and COE6 mutant n = 24) **E:** Representative images of transgenic transcriptional *unc-3* reporter [unc-3_558bp::mCherry] with wild-type COE1 or mutant COE1 sequences at L4. Top: Schematic of nerve cord section displayed in images. Arrowheads: motor neurons. Scale bar: 40 µm **F:** Quantification of average nerve cord motor neuron fluorescence intensity of transgenic transcriptional *unc-3* reporter with wild-type COE1 or mutant COE1 sequences at L4. Dots represent average nerve cord fluorescence intensities of one worm. Red bar represents median. Welch’s T-test. **G:** Model of direct, transcriptional negative autoregulation occurring at the *unc-3* endogenous locus, viewed in a gene regulatory network fashion (above) and in a molecular mechanistic model (below). For all quantifications, p-values: * < .05, ** <.01, *** < .001, n.s. = no significance.

Within the UNC-3 ChIP-seq peaks, we bioinformatically identified nine UNC-3/EBF cognate binding sites, termed COE1-9 (**Fig. 3A-B**) (see Methods). To test their functionality, we utilized CRISPR/Cas9 and substituted four nucleotides within the COE1 site (**Fig. 3A**). These exact nucleotides are necessary for transcriptional activity of UNC-3 targets in *C. elegans* MNs ^59,60^, and are likely recognized by UNC-3 based on biochemical and structural data on its mouse orthologue (EBF1) ^73,74^. Of note, the DNA binding domain (DBD) of UNC-3 shares >82% sequence similarity with the DBDs of both mouse EBF1 and human EBF3 (**Fig. 4A**). Mutation of UNC-3 binding site #1 (COE1), located in the promoter, in the context of our endogenous *unc-3::SL2::NLS::eGFP::PEST* reporter, resulted in increased eGFP expression in cholinergic MNs (**Fig. 3C-D**). Next, we tested the function of COE1 outside its genomic context. First, we generated an *mCherry* transcriptional reporter containing *unc-3*’s proximal promoter sequence (558p) that includes the COE1 site (*Punc-3_558bp::mCherry)* (**Fig. 3A**). This reporter successfully drove *mCherry* expression in ventral nerve cord MNs (**Supp. Fig. 2A-C**). Substitution of the same four nucleotides in COE1 led to increased reporter expression across multiple independent lines, suggesting that the COE1 site mediates repressive input (**Fig. 3E-F, Supp Fig. 2B-C**). Altogether, these findings indicate that *unc-3’*s negative autoregulation occurs directly and via a cognate binding site.

**Figure 4:**
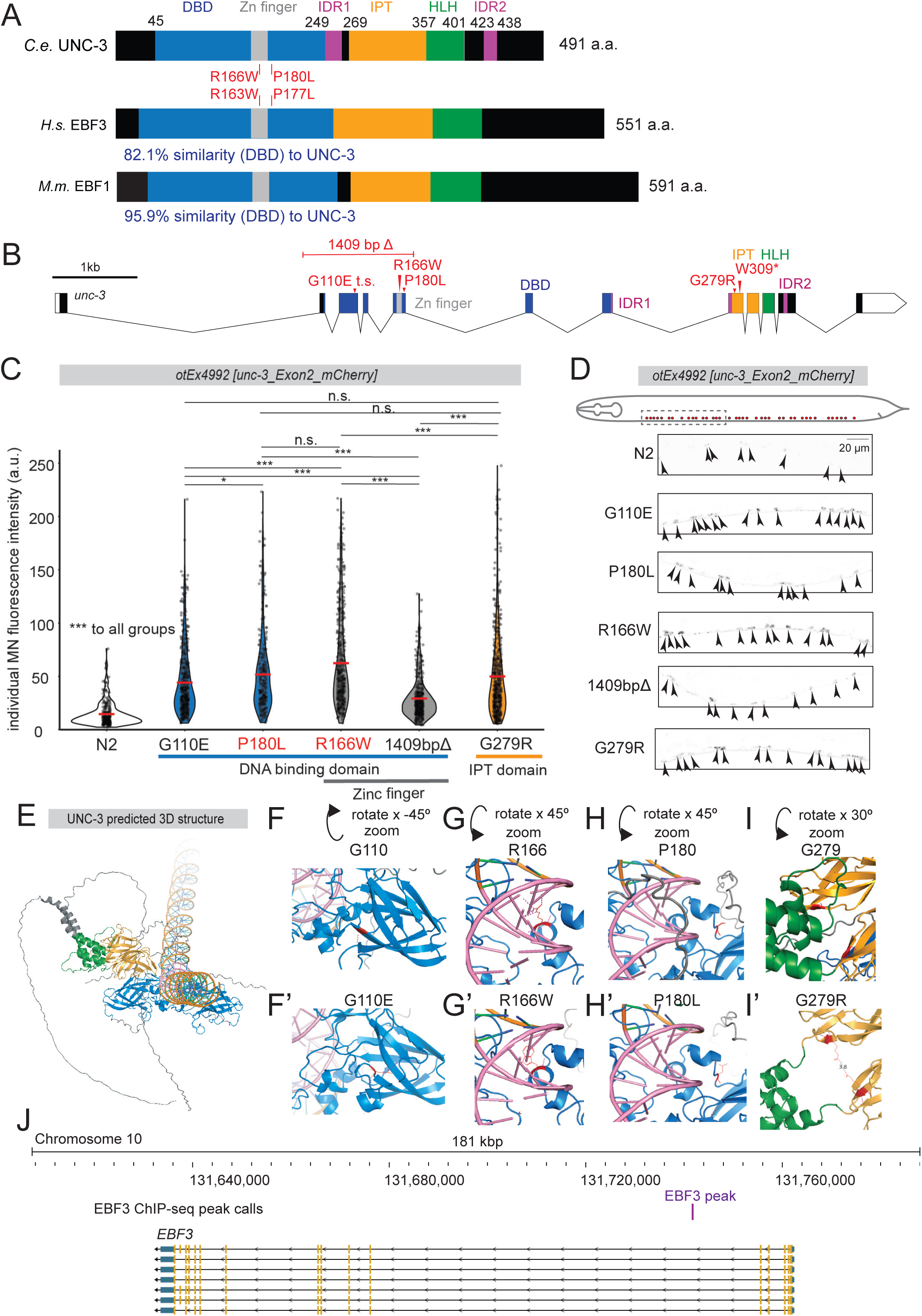
The DNA-binding and IPT domains of UNC-3 are required for self-repression. **A:** Schematics of protein domains for UNC-3 and two of its orthologues: *homo sapiens* EBF3 (hsEBF3) and *mus musculus* EBF1 (mmEBF1). Domains are colored as: blue – DNA binding domain, grey – Zinc finger, purple – Intrinsically Disordered Region (IDR), orange – Immunoglobulin/Plexin/Transcription factor (IPT) domain, green – Helix Loop Helix (HLH). Percentages show degree of similarity between DNA binding domains of EBF3 and EBF1 to DNA binding domain of UNC-3. Red mutants denote UNC-3 mutations in this study that occur in conserved residues in hsEBF3 and contribute to the EBF3-neurodevelopmental syndrome. **B:** Intron/exon schematic denoting mutants analyzed in this study (red). Protein domain color coding same as (A). Scale bar: 1 kb **C:** Quantification of individual motor neuron fluorescence intensity of fosmid-based transcriptional *unc-3* reporter at L4 in N2, *unc-3(G110E), unc-3(P180L), unc-3(R166W), unc-3(1409bpΔ),* and *unc-3(G279R)* backgrounds. Dots represent individual motor neuron fluorescence intensities. Red bar represents mean. Protein domain color coding same as (A). Dunn Kruskal-Wallis multiple comparisons test with Bonferroni correction. (N2 n = 23 worms, *unc-3(G110E)* n = 20, *unc-3 (P180L)* n = 24, *unc-3(R166W)* n = 20, *unc-3(1409bpΔ)* n = 20, *unc-3(G279R)* n = 22) **D:** Representative images of fosmid-based transcriptional *unc-3* reporter at L4 in N2, *unc-3(G110E), unc-3(P180L), unc-3(R166W), unc-3(1409bpΔ),* and *unc-3(G279R)* backgrounds. Top: Schematic of nerve cord section displayed in images. Arrowheads: motor neurons. Scale bar: 20 µm **E:** 3D structure prediction of UNC-3 homodimer bound to DNA predicted by Alphafold3. Structural domains are colored as: blue – DNA binding domain, orange – Immunoglobulin/Plexin/Transcription factor (IPT) domain, and green – Helix Loop Helix (HLH). **F-I’:** Predicted polar interactions of residues of missense mutants in WT (top) and mutated (bottom) alleles for mutants G110E **(F, F’)**, R166W **(G, G’)**, P180L **(H, H’)**, G279R **(I, I’)**. Orientation changes relative to (A) denoted above. Domain colors as shown in (A). COE binding site colored in pink. Residues of mutation shown in red. Polar contacts shown in dashed magenta line. Less than 4.0A contact shown in dashed grey line. For all quantifications, p-values: * < .05, ** <.01, *** < .001, n.s. = no significance.

### *unc-3* negative autoregulation occurs through promoter and intronic COE sites

While mutation of the COE1 site causes increased expression of the endogenous *unc-3* transcriptional reporter, it does not reach the “ceiling” of increased expression achieved when the syndromic R166W mutation is introduced (**Fig. 3D**). This suggests that *unc-3* self-repression must be occurring through additional binding sites, and/or through additional mechanisms. To test whether the additional COE sites are also involved, we used CRISPR/Cas9 to mutate COE5, located in the first intron and overlapping with the largest UNC-3 ChIP-seq peak (**Fig. 3A**). Mutation of COE5 led to increased expression of the endogenous *unc-3* transcriptional reporter, comparable to mutation of COE1 (**Fig. 3C-D**). To determine if there is a combinatorial effect, we generated a COE1 COE5 double mutant. However, the observed increase in expression was comparable to that with the mutation of either site in isolation (**Fig 3. C-D**). Further, mutation of COE site 6 also increased *unc-3* reporter expression (**Fig 3. C-D**). Taken together, these data suggest that *unc-3’*s negative autoregulation occurs directly through COE sites within its promoter and first two introns (**Fig. 3G**). These findings are highly relevant for *C. elegans* locomotion as *unc-3* movement defects can be restored only by providing *unc-3* cDNA under the control of its own promoter and first three introns ^75^. We note that while additional sites or mechanisms of *unc-3* repression remain to be identified, mutation of COE2 or COE3 sites in the background of the COE1 and 5 mutations abrogated the increase in *unc-3* expression observed in COE1 and 5 alone, as did mutation of COE8 and COE9 sites in the background of the COE6 mutation (**Supp. Fig. 2D-F**).

### Mutations in DBD, Zn finger, and IPT domains disrupt *unc-3* negative autoregulation

To gain mechanistic insight, we sought to determine which protein domains are necessary for *unc-3* self-repression. The UNC-3 protein contains several highly conserved domains (**Fig. 4A**): a Zn-finger-containing DBD involved in DNA binding site recognition ^76^; an Immunoglobulin/Plexin/Transcription Factor (IPT) domain of uncertain function; and a Helix-Loop-Helix (HLH) domain associated with homo-dimerization ^73,76,77^. Further, we bioinformatically identified two Intrinsically Disordered Regions (IDRs): one in the linker region between the DBD and IPT domain (IDR1), and one in the C-terminus (IDR2) (**Fig. 4A**) (see Methods).

To determine the necessity of each domain in *unc-3* negative autoregulation, we first analyzed strains with missense mutations in the DBD (G110E, P180L), Zn finger (R166W), or IPT domains (G279R) (**Fig. 4B**). As a positive control, we used a strong LoF mutation consisting of an in-frame deletion of 1,409 bp that removes the entire Zn finger and a portion of the DBD (**Fig. 4B**). All mutations led to increased expression of the *unc-3_Exon2_mCherry* reporter in cholinergic MNs. The mutants produced an allelic series (N2 < *1409 bp deletion* < *G110E, P180L, G279R* < *R166W*), with the strongest effect observed with the syndromic R166W mutation that specifically affects the Zinc finger domain (**Fig. 4C-D**).

To predict at the structural level how each missense mutation may disrupt UNC-3 function, we generated AlphaFold predictions (see Methods) (**Fig. 4E-I’**). We analyzed putative changes in polar contacts (as a proxy for hydrogen bonds), which are significant contributors to protein structure and TF-DNA interaction specificity ^78^. For DBD mutations, different effects were predicted: G110E disrupts DBD structure (**Fig. 4F-F’**); R166W disrupts interaction between the DBD and the COE site in DNA (**Fig. 4G-G’**); and P180L showed no immediate polar contact changes (**Fig. H-H’**). The mutation within the IPT, G279R, showed no polar contact changes, but did demonstrate a new contact of less than 4.0A between IPT domains of the two UNC-3 monomers, potentially influencing UNC-3 homodimerization (**Fig. I-I’**). Overall, these findings suggest that the DBD, Zinc finger, and IPT domains of UNC-3 are important for its negative autoregulation.

### *EBF3* syndromic mutations disrupt *unc-3* negative autoregulation

Both R166W and P180L mutations are found in human EBF3 and cause the HADDS neurodevelopmental syndrome ^67^ (**Fig. 4A**). We found that both mutations increase *unc-3* reporter expression in *C. elegans* MNs (**Fig. 4C-D**). To determine whether the two additional mutations (G110E, G279R) that increase *unc-3* expression could have clinical relevance, we interrogated the AlphaFold Missense dataset, which assigns a pathogenicity score for every single amino acid substitution in the human protein-coding genome ^79,80^. Both missense mutants (G110E, G279R) are predicted to be pathogenic (**Supp. Table 1**). In addition, ChIP-seq data of human EBF3 ^57^ demonstrates that EBF3 binds at its own locus (**Fig. 4J**). Altogether, our data on four missense mutations (R166W, P180L, G110E, G279R) suggest disruption of *EBF3* negative autoregulation as a potential pathogenetic mechanism in HADDS.

### The UNC-3 DBD is both necessary and sufficient for *unc-3* negative autoregulation

The missense mutations described above do not target every protein domain of UNC-3 (e.g., HLH, IDR) (**Fig. 4B**). To more comprehensively assess the function of each domain in *unc-3* negative autoregulation, we conducted a rescue assay. We reintroduced the UNC-3 protein (cDNA) with varying protein domains removed (**Fig. 5A**) into *unc-3*(W309*) LoF (presumptive null) animals that also carried the transgenic *unc-3_Exon2_mCherry* reporter (**Fig. 5B**). Reintroduction of wild-type *unc-3* cDNA under the control of the *unc-3* promoter (558bp) rescued the loss of expression of known UNC-3 target genes (*cho-1/ChT* and *unc-129/TGFB)* ^59^ (**Supp. Fig. 3D,H**) and thrashing speed defects (**Fig. 5K**), suggesting this promoter is driving UNC-3 expression in a functionally relevant manner. Reintroduction of the wild-type *unc-3* cDNA also led to a significant, albeit partial, reduction of the increased *unc-3* reporter expression observed in *unc-3(*W309*) mutants when compared to isogenic controls (**Fig. 5B-C**). The partial effect is in part due to the *unc-3* promoter fragment (558bp) being active in most, but not all, of the 39 cholinergic MNs (**Supp. Fig. 2A**). Reintroduction of UNC-3 lacking the DBD (ΔDBD) did not rescue the increased *unc-3* reporter expression (**Fig. 5D**), consistent with the effect of the DBD missense mutations (G110E, P180L, R166W) (**Fig. 4C-D**). The ΔDBD construct also failed to rescue thrashing speed defects (**Fig. 5K**), corroborating a requirement for the DBD in both *unc-3*’s negative autoregulation and locomotory behavior.

**Figure 5:**
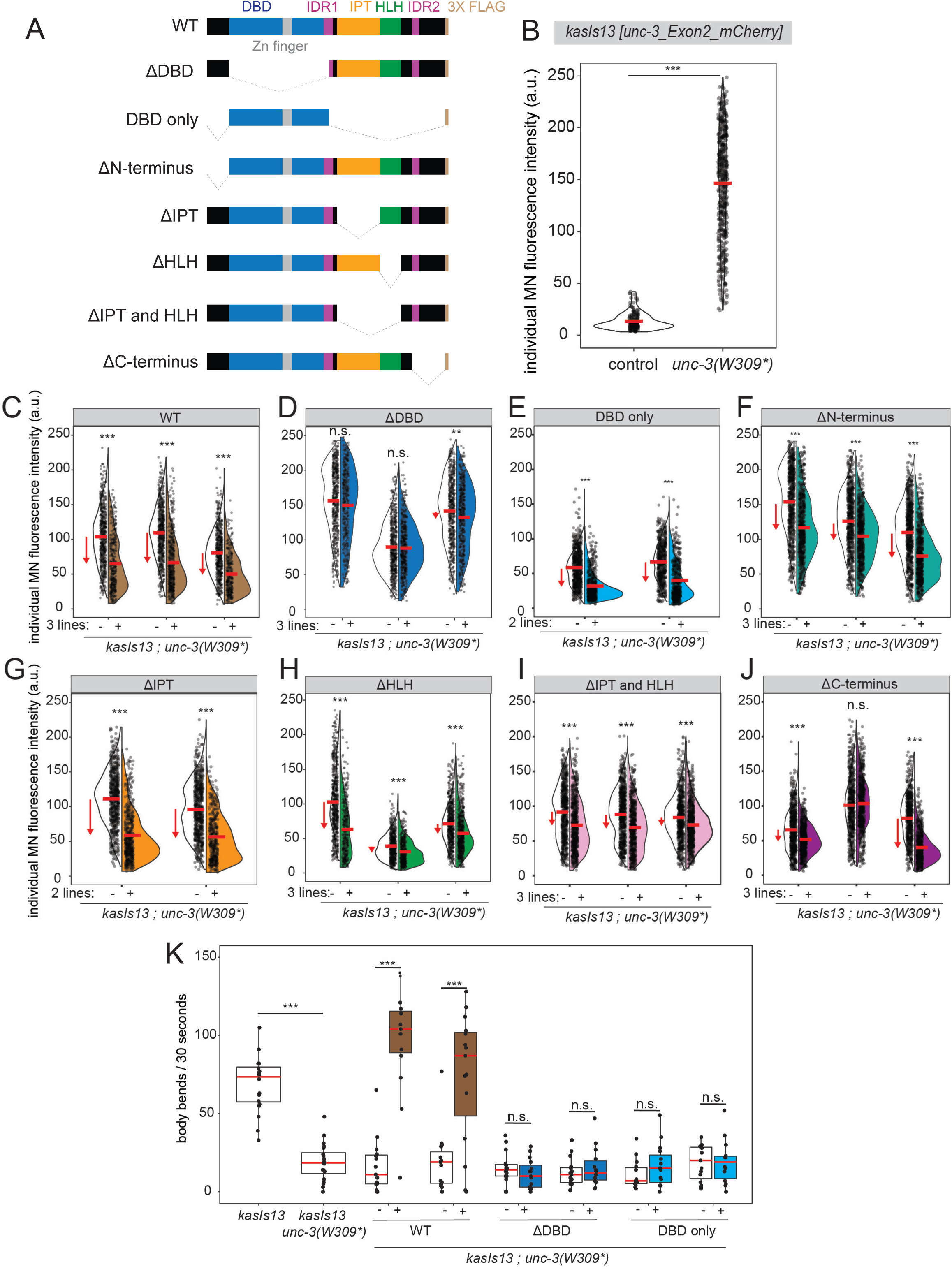
The DNA-binding domain of UNC-3 is necessary and sufficient for self-repression. **A:** Schematic of cDNA constructs driven by the 558bp promoter of *unc-3* in the rescue assay. **B:** Quantification of individual motor neuron fluorescence intensity of fosmid-based transcriptional *unc-3* reporter at L4 in control and *unc-3(W309*)* backgrounds. Dots represent individual motor neuron fluorescence intensities. Red bar represents mean. Welch’s T-test. (control n = 8 worms, *unc-3(W309*)* n = 15) **C-J:** Quantification of individual motor neuron fluorescence intensity of fosmid-based transcriptional *unc-3* reporter at L4 in *unc-3(W309*)* background either with (right, colored side of split violin) or without (left, uncolored side of split violin) the respective rescue construct (top). Dots represent individual motor neuron fluorescence intensities. Red bar represents mean. Red arrow highlights mean change in construct-carrying vs non-construct-carrying siblings. Construct consists of Punc-3_558bp driving unc-3 cDNA **(C)** WT (kasEx356+ n = 19 worms, kasEx356- n = 20, kasEx358+ n = 21, kasEx358- n = 20, kasEx357+ n = 20, kasEx357- n = 20) **(D)** DNA binding domain (DBD) deletion (kasEx364+ n = 20, kasEx364- n = 19, kasEx363+ n = 20, kasEx363- n = 20, kasEx362+ n = 19, kasEx362-n = 20) **(E)** only DBD (kasEx376+ n = 17, kasEx376- n = 22, kasEx377+ n = 16, kasEx377- n = 19) **(F)** N-terminal deletion (kasEx359+ n = 10 worms, kasEx359- n = 19, kasEx360+ n = 20, kasEx360- n = 20, kasEx361+ n = 21, kasEx361- n = 21), **(G)** Immunoglobulin/Plexin/Transcription factor (IPT) domain deletion (kasEx365+ n = 20, kasEx365- n = 20, kasEx366+ n = 20, kasEx366- n = 21), **(H)** Helix Loop Helix (HLH) deletion (kasEx368+ n = 21, kasEx368- n = 22, kasEx367+ n = 20, kasEx367- n = 20, kasEx369+ n = 20, kasEx369- n = 20), **(I)** HLH and IPT domain deletion (kasEx370+ n = 20, kasEx370- n = 19, kasEx362+ n = 21, asEx362- n = 20, kasEx371+ n = 19, asEx371- n = 19), **(J)** C-terminus deletion (kasEx375+ n = 20, kasEx375- n = 21, kasEx373+ n = 19, kasEx373- n = 20, kasEx374+ n = 19, kasEx374- n = 20). Welch’s T-test. **K:** Quantification of body bends per 30 seconds of control (*kasIs13*), mutant *(kasIs3 unc-3(W309*)*, and array-carrying worms (brown – WT rescue; dark blue – DBD deletion; bright blue – DBD only) with isogenic sibling controls (white). (kasIs13 n = 20 worms, kasIs13 *unc-3(W309*)* n = 20), WT (kasEx356- n = 15 worms, kasEx356+ n = 15, kasEx358- n = 15, kasEx358+ n = 15), DBD deletion (kasEx364- n = 15 worms, kasEx364+ n = 15, kasEx362- n = 15, kasEx362+ n = 14), DBD only (kasEx376- n = 14 worms, kasEx376+ n = 15, kasEx377- n = 15, kasEx377+ n = 14). Each dot represents one worm. Red bar represents mean. Welch’s T-test. (kasIs13 n = 20 worms, kasIs13 *unc-3(W309*)* n = 20). For all quantifications, p-values: * < .05, ** <.01, *** < .001, n.s. = no significance.

To test whether the DBD alone is sufficient for *unc-3* self-repression, we expressed only the DBD in *unc-3* (W309*) LoF animals. Strikingly, introduction of the DBD alone still rescued *unc-3* reporter repression (**Fig. 5E**); however, it failed to rescue thrashing speed defects (**Fig. 5K**). Overall, these data demonstrate that the DBD is both necessary and sufficient to drive *unc-3* negative autoregulation, while additional domains are required for UNC-3-dependent motor behavior (thrashing).

Interestingly, rescue of increased *unc-3* expression still occurred even upon individual deletion of either the N-terminus (ΔN-terminus), IPT domain (ΔIPT), HLH domain (ΔHLH), or IDR-containing C-terminus (ΔC-terminus), suggesting that these domains are dispensable for UNC-3 repressive capacity in the context of this rescue assay (**Fig. 5F-H, J**). To test for potential functional redundancy between the IPT and HLH domains, we deleted them both simultaneously (ΔIPT and HLH); while a reduction of *unc-3* reporter expression still occurred, it was to a lesser magnitude as compared to the full-length UNC-3 or individual IPT and HLH deletions (**Fig. 5I**), suggesting partial cooperative contributions of these domains to optimal repressive activity.

### The DBD and IPT domains are also necessary for activation of UNC-3 target genes

Prior work has focused on the function of UNC-3 as a transcriptional activator of multiple effector genes in cholinergic MNs^58–62,81^. This is in sharp contrast with *unc-3* directly repressing its own expression. How does UNC-3 act as both a direct self-repressor and activator of effector genes in cholinergic MNs? To address this question, we first assessed whether the domains involved in *unc-3* self-repression were also important for UNC-3’s function as a transcriptional activator. We crossed the same *unc-3* mutant alleles (G110E, P180L, R166W, 1409bp deletion, G279R) to a reporter strain for *cho-1/ChT,* a target gene directly activated by UNC-3 ^59^. In all mutants, *cho-1/ChT* reporter expression decreased (**Supp. Fig. 3A-B**), suggesting that the DBD, Zinc finger, and IPT domains of UNC-3 are important for activating gene expression in addition to their necessity for *unc-3* repression.

These findings suggest that the missense mutants available are not sufficient to genetically dissect between the repressor and activator functions of UNC-3, potentially because the same domain(s) contribute to both self-repression and activation. We therefore surveyed additional domains using the previously described rescue assay, this time assessing rescue of activation of *cho-1/ChT* expression in *unc-3* (W309*) LoF animals (**Supp. Fig. 3C**). Reintroduction of wild-type UNC-3 restored *cho-1* reporter expression relative to isogenic controls (**Supp. Fig. 3D**). Deletion of the HLH and IPT domain in combination (ΔIPT and HLH), or of the C-terminus (ΔC-terminus), also restored *cho-1* expression, suggesting that these domains are dispensable for UNC-3’s activator function in the context of this rescue assay (**Supp. Fig. 3F-G**). Consistent with the missense mutant analysis (**Supp. Fig. 3A-B**), deletion of the DBD (ΔDBD) abrogated rescue of *cho-1/ChT* expression (**Supp. Fig. 3E**), corroborating its essential role for UNC-3-mediated gene activation.

To determine whether the DBD was sufficient to mediate UNC-3 activating function, we assessed rescue of expression of another directly activated target, *unc-129/TGFB*, in *unc-3(*W309*) LoF animals. Introduction of wild-type UNC-3 rescued *unc-129* reporter expression relative to isogenic controls to a level of expression comparable to the reporter in a wild-type background (**Supp. Fig. 3H**). Introduction of only the DBD led to a partial rescue of *unc-129* expression (**Supp. Fig. 3H**), suggesting that the DBD alone possess limited intrinsic activating capacity, but that additional domains are required for complete transcriptional activation efficiency.

### An *in silico* AlphaFold2 screen identifies transcription and chromatin factors as putative UNC-3-interacting cofactors

Given that UNC-3’s DBD and IPT domains are necessary for both activation and repression, we then sought to determine how these domains may contribute to these functions. As the DBD is both necessary and sufficient for *unc-3’s* self-repression, one hypothesis is that the DBD limits *unc-3’s* transcriptional activation by preventing other TFs (e.g., steric hindrance) from binding to the *unc-3* locus; however, we found no overlap observed between COE sites and the binding sites of known *unc-3* activators (Hox/PBX) (**Supp. Fig. 4A**). An additional mechanism could be that the DBD functions to recruit corepressors. To identify such factors, we conducted an *in silico* protein-protein interaction screen using LocalColabFold, an implementation AlphaFold2 (see Methods) ^82,83^. We tested for pairwise interactions between UNC-3 and all of the TFs and chromatin factors known to be expressed in nerve cord MNs ^84^. Candidates were selected based on a previously used prediction quality cut-off of at least a 0.3 interface predicted template modeling (ipTM) score for the majority (3 out of 5) of the models^63,85^ (**Supp. Fig. B-C**). Further refinement through literature search resulted in a list of 11 putative interactors (**Supp. Fig. 4D**). Three TFs (HLH-8/TWIST, MXL-2/MLX, MXL-1/MAX) are predicted to interact via the HLH and/or IPT domains of UNC-3. Notably, the remaining eight proteins are chromatin factors predicted to interact with UNC-3’s DBD (MTQ-2/HEMK2, HDA-3/HDAC1-2, HDA-6/HDAC6, T22B7.4/SGF29, Y17G9B.8/SGF29, TRR-1/TRRAP, ISW- 1/SMARCA1) or IPT domain (SPR-5/KDM1A). In summary, this *in silico* screen suggests that UNC-3 may modulate its activator and repressor activities through recruitment of TFs via its HLH/IPT domains, or of chromatin factors via its DBD or IPT domains.

### Motor neuron terminal identity is sensitive to UNC-3 levels

Negative autoregulation of *unc-3* suggests that it is critical to regulate UNC-3 levels within a specific range. If this were correct, we would predict that over- or under-expression of *unc-3* will affect MN molecular identity. To test this, we assessed expression of two terminal identity genes (*cho-1/ChT* and *unc-129/TGFB)* (**Supp. Fig. 5A**) upon UNC-3 over-expression using two different cholinergic MN-specific promoters. Over-expression of UNC-3 driven by the *ace-2* promoter ^59^ (*Pace-2::unc-3_cDNA*) caused increased *unc-129::GFP* expression in cholinergic MNs (**Supp. Fig. 5B-C**). Similarly, over-expression of UNC-3 with the *unc-3* promoter (*Punc-3::unc-3_cDNA*) led to increased expression of both *unc-129::GFP* and *cho-1/ChT* (**Supp. Fig. 5D-E**). Next, we sought to determine whether MN identity was affected upon decreasing *unc-3* dosage. In heterozygous and homozygous animals for the LoF allele *unc-*3(W309*), both *unc-129::GFP* and *cho-1::mChOpti* expression decreased in a dosage-dependent manner (**Fig. 6A-B**), indicating haploinsufficiency. Altogether, these data suggest that MN terminal identity is sensitive to UNC-3 levels.

**Figure 6:**
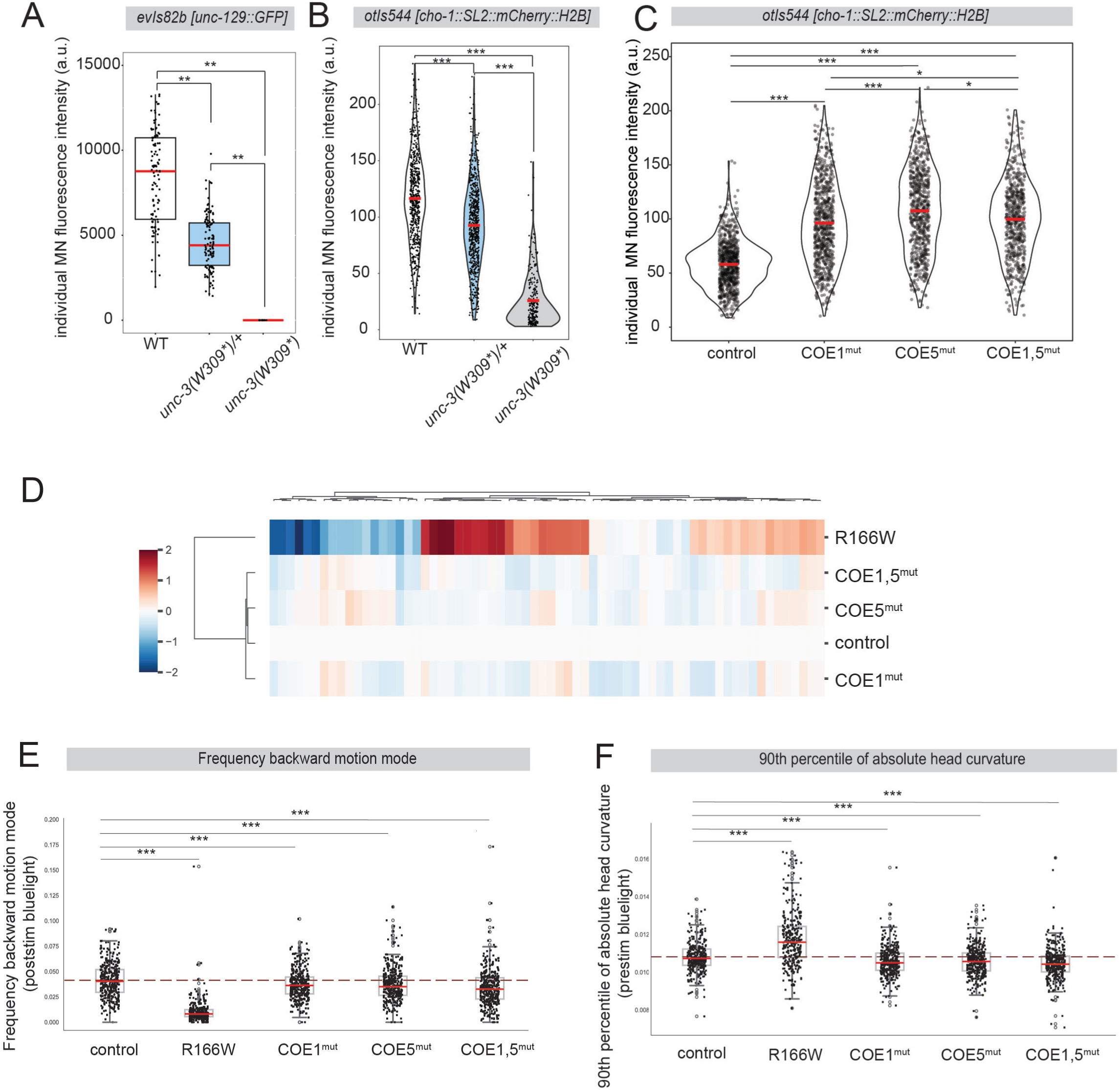
Disruption of *unc-3*’s negative autoregulation affects MN identity and animal locomotion. **A:** Quantification of individual motor neuron fluorescence intensity of transgenic *unc-129* reporter at L4 in N2, *unc-3(W309*/+)*, or *unc-3(W309*)* backgrounds. Dots represent individual motor neuron fluorescence intensities. Red bar represents mean. Tukey honest significance test. (N2 n = 13 worms, *unc-3(W309*/+)* n = 12, *unc-3(W309)* n = 11) **B:** Quantification of individual motor neuron fluorescence intensity of fosmid-based transcriptional *cho-1* reporter at L4 in N2, *unc-3(W309*/+)*, or *unc-3(W309*)* backgrounds. Dots represent individual motor neuron fluorescence intensities. Red bar represents mean. Tukey honest significance test. (N2 n = 16, *unc-3(W309*/+)* n = 19, *unc-3(W309)* n = 20) **C:** Quantification of individual motor neuron fluorescence intensity of fosmid-based transcriptional *cho-1* reporter at L4 in control, COE1mut, COE5mut, and COE1,5mut. Dots represent individual motor neuron fluorescence intensities. Red bar represents mean. Dunn Kruskal-Wallis multiple comparisons test with Bonferroni correction. **D:** Hierarchically clustered heatmap of 22-feature analysis of automated worm tracking behavior. **E:** Quantification of frequency of worm backwards motion after stimulation with blue light. Welch’s T-test to control for each group. Outliers shown in open circles. Solid red bar represents median. Dashed red bar represents mean of the control. **F:** Quantification of 90^th^ percentile of absolutely head curvature prior to stimulation with blue light. Welch’s T-test to control for each group. Outliers shown in open circles. Solid red bar represents median. Dashed red bar represents mean of the control. For all quantifications, p-values: * < .05, ** <.01, *** < .001, n.s. = no significance.

### Minimal disruption of *unc-3*’s negative autoregulation leads to MN and locomotion defects

Given the importance of *unc-3* dosage in MN terminal identity, we predicted that increasing *unc-3* levels by disrupting negative autoregulation would affect MN gene expression and locomotory behavior. To test this, we used three of the *cis-*regulatory *unc-3* mutants that demonstrated increased *unc-3* expression: COE1^mut^, COE5^mut^, and COE1,5^mut^ (**Fig. 3C-D**). We found that *cho-1/ChT* expression increased in all three mutants, suggesting that minimal disruption of *unc-3*’s negative autoregulation affects MN terminal identity (**Fig. 6C**).

To finely dissect the specific contributions of negative autoregulation to locomotion, we analyzed the behavior of COE1^mut^, COE5^mut^, and COE1,5^mut^ *cis*-regulatory mutants. We adopted a high-resolution phenomics approach of freely moving worms using automated multi-worm tracking, offering a quantitative analysis of locomotion (e.g., curvature, speed, forward and backward motion frequency)^86^. Minimal disruption of *unc-3* negative autoregulation through the COE1^mut^, COE5^mut^, and COE1,5^mut^ mutations (**Fig. 3C-D**) resulted in broad locomotory effects (**Fig. 6D, Supp. Fig. 5F**). Analysis of specific traits showed disruption in motion modes, wherein *cis*-regulatory mutant animals had a decrease in pausing frequency and backward frequency after blue light stimulation relative to controls (**Fig. 6E, Supp. Fig. 5G**). *Cis*-regulatory mutant animals also showed changes in posture, specifically decreases in head curvature and an increase in length (**Fig. 6F, Supp. Fig. 5H-J**), and changes in head angular velocity (**Supp. Fig. 5K**). We then assessed how these changes compared to LoF of UNC-3 in *unc-3(R166W)* animals. Hierarchical clustering showed that the LoF *unc-3(R166W)* mutation showed the most severe locomotion defects (**Fig. 6D, Supp. Fig. 5F**), as this mutation has a strong impact not only on *unc-3* levels, but also on MN terminal identity genes. *unc-3(R166W)* animals demonstrated similar behavioral effects on motion mode, but at a much greater magnitude (**Fig. 6E, Supp. Fig. 5G**). In contrast, effects on posture trended in opposite directions in *cis*-regulatory mutants relative to *unc-3(R166W)* (**Supp. Fig. 5H-J**), highlighting unique contributions of *unc-3* levels on locomotion. Altogether, these data show that minimal disruption of *unc-3*’s negative autoregulation leads to MN identity and locomotion defects.

### *unc-3*’s negative autoregulation is continuously counteracted by CEH-20/PBX

Despite its self-repression, UNC-3 is still expressed in cholinergic MNs throughout life (**Supp. Fig. 6A**), suggesting that additional factors must be providing positive input to *unc-3*, thereby counteracting negative autoregulation. We previously showed that the midbody Hox proteins LIN-39 (*Scr/Dfd/Hox3-5*) and MAB-5 (*Antp/Hox6-8*) can activate *unc-3* expression^51^.

However, simultaneous loss of both *lin-39* and *mab-5* reduces *unc-3* expression only by ∼50% in cholinergic MNs ^51^, suggesting additional activators remain to be identified. Because HOX proteins are known to function in collaboration with cofactors ^87^, we tested whether the Hox cofactor CEH-20, ortholog of the conserved Pre-B-cell leukemia (PBX) family of TFs, is required for *unc-3* expression. In animals carrying a homozygous *ceh-20(ok541)* null allele^88^ (**Fig. 7A**), we found a striking decrease in the expression of both a transcriptional *unc-3* reporter (*Punc-3_223bp::mCherry*) and an endogenous UNC-3 translational reporter (UNC-3::GFP) at L2 (**Fig. 7B-D**). In *ceh-20* hypomorphic mutants (*ceh-20(ay42)*)^89^ (**Fig. 7A**), we also found decreased expression of UNC-3::GFP both at L2 and L4 (**Fig. 7E**). Available ChIP-seq data for CEH-20 at L4 demonstrate binding at the *unc-3* locus, supporting a mechanism of direct activation (**Fig. 7B**) ^90^. Taken together, these data suggest that CEH-20/PBX directly activates UNC-3 expression in cholinergic MNs.

**Figure 7:**
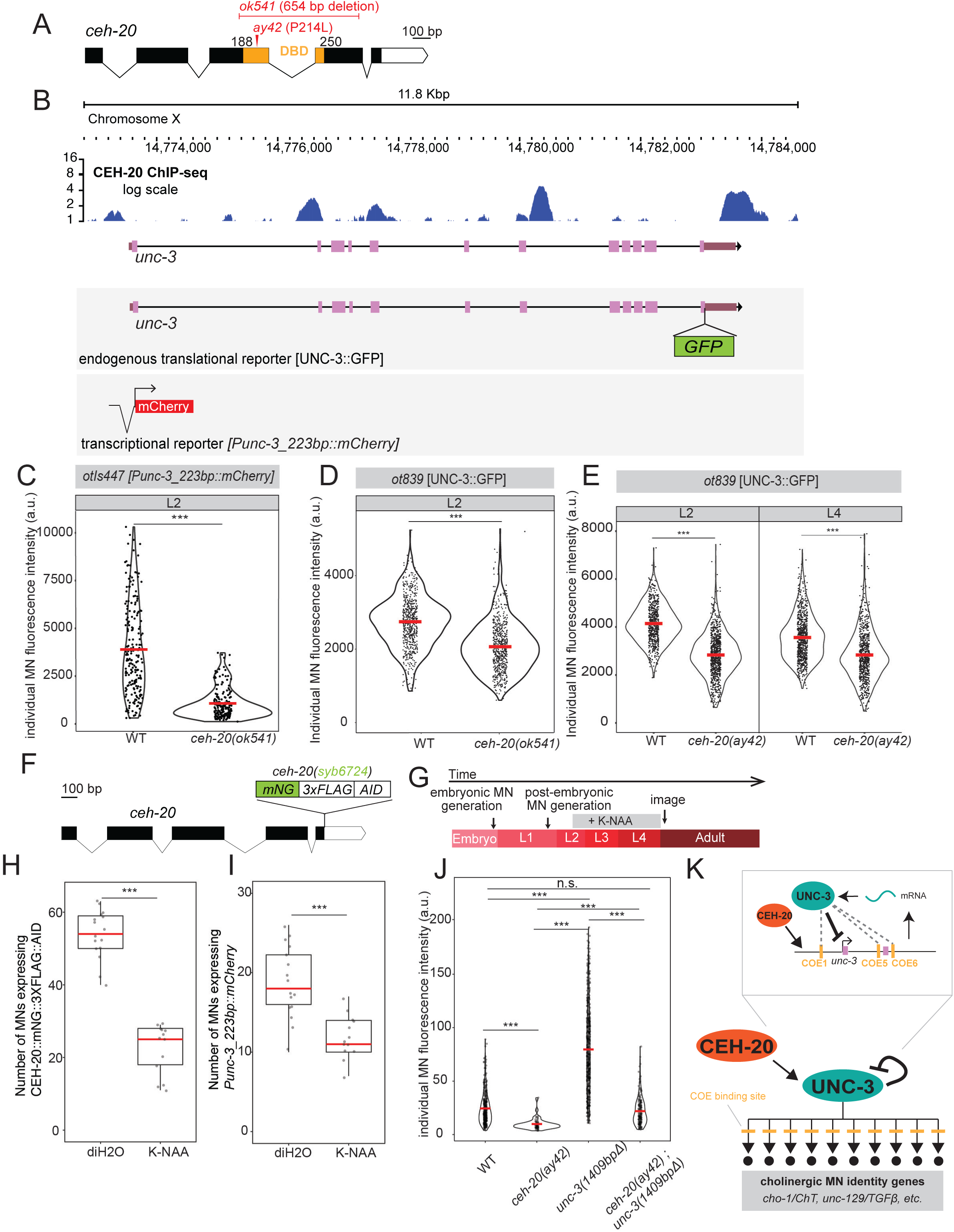
Continuous positive input by CEH-20/PBX on *unc-3* expression. **A:** Intron/exon schematic of *ceh-20* gene with mutants used in this study marked in red. Orange – DNA binding domain (DBD). Scale bar: 100 bp **B:** Genomic view of the endogenous *unc-3* locus with CEH-20 ChIP-seq tracks, log scale (whole animal, L4, fold change over control). Below *unc-3* locus: molecular nature of *unc-3* endogenous translational reporter, and transgenic transcriptional reporter. **C:** Quantification of individual motor neuron fluorescence intensity of transgenic transcriptional *unc-3* reporter at L2 in N2 or *ceh-20(ok541)* backgrounds. Dots represent individual motor neuron fluorescence intensities. Red bar represents mean. Welch’s T-test. (N2 n = 9, *ceh-20(ok541)* n = 18) **D:** Quantification of individual motor neuron fluorescence intensity of *unc-3* endogenous translational reporter at L2 in N2 or *ceh-20(ok541)* backgrounds. Dots represent individual motor neuron fluorescence intensities. Red bar represents mean. Welch’s T-test. (N2 n = 16, *ceh-20(ok541)* n = 21) **E:** Quantification of individual motor neuron fluorescence intensity of *unc-3* endogenous translational reporter at L2 (left) and L4 (right) in N2 or *ceh-20(ay42)* backgrounds. Dots represent individual motor neuron fluorescence intensities. Red bar represents mean. Welch’s T-test (L2: N2 n = 16, *ceh-20(ay42)* n = 22, L4: N2 n = 20, *ceh-20(ay42)* n = 20) **F:** Intron/exon schematic of *ceh-20* gene with endogenous tags of mNeonGreen (mNG), 3XFLAG, and auxin inducible degron (AID). **G:** Schematic of timing of auxin exposure and imaging. **H:** Quantification of number of motor neurons expressing endogenous CEH-20::mNG::3XFLAG::AID at day 1 adulthood in control (diH2O) or auxin conditions. Welch’s T-test. (diH2O n = 16 worms, auxin n = 13). **I:** Quantification of number of motor neurons expressing transcriptional *Punc-3_223bp::mCherry* reporter at day 1 adulthood in control (diH2O) or auxin conditions. Welch’s T-test. (diH2O n = 16 worms, auxin n = 13). **J:** Quantification of individual motor neuron fluorescence intensity of fosmid-based transcriptional *unc-3* reporter at L4 in N2, *ceh-20(ay42), unc-3(n3435),* or *ceh-20(ay42) ; unc-3(n3435)* backgrounds. Dots represent individual motor neuron fluorescence intensities. Red bar represents mean. Dunn Kruskal-Wallis multiple comparisons test with Bonferroni correction (N2 n = 19 worms, *ceh-20(ay42)* n = 20, *unc-3(1409bpΔ)* n = 20, *ceh-20(ay42) ; unc-3(1409bpΔ)* n = 20) **K:** Schematics representing the ‘balancing-act’ model of converging activating and repressive inputs at the *unc-3* locus, viewed in a gene regulatory network fashion (above) and in a molecular mechanistic model (below). For all quantifications, p-values: * < .05, ** <.01, *** < .001, n.s. = no significance.

To determine whether CEH-20 functions to maintain UNC-3 expression, we again used the AID system, this time using a *ceh-20::mNG::3XFLAG::AID* allele (**Fig. 7F**) ^91^. Continuous auxin exposure from the end of L2 stage until the first day (day 1) of adulthood led to depletion of CEH-20 (**Fig. 7G-H**). This was accompanied by a decrease in the total number of cells expressing an *unc-3* transcriptional reporter (**Fig. 7I**), suggesting that CEH-20 is required to maintain activation of *unc-3* expression after L2.

Is CEH-20 functioning in opposition of UNC-3 to regulate *unc-3* levels? To assess this, we generated *ceh-20(ay42); unc-3(1409 bp deletion)* double mutant animals. Consistent with previous experiments (**Fig. 7C-E**, **Fig. 4C**), single *ceh-20(ay42)* mutants display a decrease in *unc-3_Exon2_mCherry* reporter expression in MNs, whereas *unc-3(1409bp deletion)* animals show an increase (**Fig. 7J**). However, double mutants display significantly lower levels of *unc-3* expression compared to *unc-3(1409bp deletion*) mutant animals (**Fig. 7J**), suggesting that UNC-3 negative autoregulation counteracts the activating function of CEH-20 on the *unc-3* locus (**Fig. 7K**).

## DISCUSSION

We describe here a gene regulatory mechanism that continuously fine-tunes expression of a dosage-sensitive and clinically relevant TF in *C. elegans* motor neurons. Our study provides the first example of negative autoregulation of a terminal selector gene, enriching our understanding of how these critical regulators of neuronal identity and function maintain their expression at precise levels in long-lived, post-mitotic cells. Because deletions, duplications, and enhancer mutations in *EBF3* (human *unc-3* ortholog) alter its expression levels and cause a severe neurodevelopmental syndrome (HADDS) ^7,8^, our findings of HADDS patient variants disrupting *unc-3* negative autoregulation suggest a putative pathogenic mechanism centered on dysregulated *EBF3* levels.

Negative autoregulation of TFs has been mainly studied in synthetic, prokaryotic, and developmental contexts, where it can have diverse spatiotemporal effects on gene expression^19,20^, such as generation of oscillations^92^, or restriction of expression variability ^30^. However, the importance of negative autoregulation in maintenance of long-lived cell types has not yet been investigated. Using genome engineering, we show that minimal disruption of *unc-3*’s negative autoregulation leads to MN identity and locomotion defects in *C. elegans*. Mechanistically, TF negative autoregulation can occur either at the transcriptional (e.g., *Drosophila* temporal TFs^93^), or post-transcriptional (e.g., PITX3 in dopaminergic neurons^94^) level. In this study, we reveal a case of transcriptional negative autoregulation that occurs directly through cognate UNC-3/EBF sites (TCCCNNGGGA) found in the promoter and first two introns of *unc-3*. Interestingly, *unc-3* movement defects can be restored only by providing *unc-3* cDNA under the control of its own promoter and first three introns^75^, corroborating a physiological role for negative autoregulation in maintaining *unc-3* expression at precise levels. Notably, we were unable to reach the “ceiling” of *unc-3* expression increase through our mutational analysis of cognate UNC-3/EBF (COE) sites, suggesting that additional mechanisms are needed to mediate *unc-3* repression. These may involve non-canonical UNC-3/EBF sites ^58^, or indirect negative autoregulation through repression of *unc-3* activators, such as LIN-39/HOX^51^. Furthermore, the identification of COE sites, which upon mutation abrogate the increased *unc-3* expression seen in COE1, COE5, or COE6 regulatory mutants (**Supp. Fig. 2D-F**), suggests distinct COE sites may be required for either positive or negative autoregulation. These counteracting autoregulatory inputs could occur simultaneously throughout life, in a “tug-of-war” type of mechanism; alternatively, *unc-3* may positively autoregulate through certain COE sites early in MN development, but later switch to negative autoregulation through different COE sites to avoid run-off amplification.

At the protein level, we demonstrate that negative autoregulation of UNC-3 is mediated by its DBD and IPT domain. The sufficiency of the DBD to repress *unc-3* suggests that this domain is a major contributor to *unc-3’s* repressive capacity. While the mechanism remains unclear, two non-mutually exclusive scenarios could be envisioned. First, the DBD could function to sterically hinder the binding or activity of *unc-3* activators (e.g., LIN-39/HOX, CEH-20/PBX) ^95^. However, our *cis*-regulatory analysis of binding sites for UNC-3/EBF and HOX or PBX on the *unc-3* locus did not reveal any overlap, arguing against steric hindrance. Second, the DBD could mediate protein-protein interactions with chromatin factors or TFs, resulting in *unc-3* repression. Supporting this possibility, our *in silico* screen identified multiple chromatin factors capable of repression that are predicted to interact specifically with the DBD of UNC-3. Outside the ventral nerve cord, UNC-3 has previously been found to directly interact with the H3K27me2/3 demethylase JMJD-3.1/JMJD3 and the histone acetyltransferase CBP-1/P300 ^96,97^. However, these chromatin factors are generally associated with gene activation, and the UNC-3 domain mediating the interaction remains unknown.

Although the function of UNC-3 as a transcriptional activator in *C. elegans* MNs is well established^59–62,81^, whether and how it is involved in gene repression is poorly understood. By discovering *unc-3*’s negative autoregulation, we offer mechanistic insights into UNC-3-mediated gene repression possibly applicable to other genes repressed by UNC-3. Of note, our recent work identified ∼1,200 genes repressed by UNC-3 in MNs, but the underlying mechanisms remain unknown^58^.

Which mechanisms dictate UNC-3’s function as a transcriptional activator versus repressor? UNC-3 acts directly through canonical UNC-3/EBF (COE) sites not only to activate its targets^59^, but also to self-repress (this study), suggesting that the functional dichotomy of UNC-3 depends on its protein domains. Here, we found both the activating and self-repressing functions require its DBD and IPT domains. Unlike the IPT domain of other TFs (e.g., NF-kB) that contributes to both DNA contact and dimerization, the IPT of the UNC-3/EBF family is suggested to only contribute to dimerization^73^. Therefore, the ability of UNC-3 to dimerize via the IPT, either with itself or other factors, appears necessary for overall UNC-3 function. Furthermore, our protein deletion rescue assay did not identify additional domains required for either self-repression or activation, including the IDR-containing C-terminus. However, it did reveal that the UNC-3 DBD by itself can function as a weak transcriptional activator, consistent with prior work showing that the DBD of human EBF1 has modest activating function^76^ . Despite this, the DBD was not sufficient to rescue worm swimming speeds, suggesting additional domains are necessary for UNC-3-mediated transcription and proper motor behavior. Overall, these findings suggest that UNC-3’s function as both an activator and a repressor depends on its DNA binding capacity, and the separation of these functions may be determined by the protein-protein interactions mediated by the DBD and/or IPT domain. Last, our protein domain assays in MNs may not only illuminate the dual function of UNC-3 in other *C. elegans* neuron types ^98^, but also of *EBF3* in human neurons.

A priori, a repressive input, such as negative autoregulation, is not necessarily required to dampen TF expression levels. For example, a TF may not have a dosage-sensitive function, or expression could be dampened through other mechanisms (e.g., weak activating input, fast protein turnover, slow rate of transcription) ^99,100^. However, we show that a continuous activating input provided by HOX/PBX proteins is counteracted by *unc-3* negative autoregulation. Why combine a continuous positive input with continuous negative autoregulation? Negative autoregulation can function to restrict TF expression levels within a critical range ^30^, an important function given that variation in TF expression can disrupt development and contribute to disease ^6^. This has been well demonstrated in *Drosophila,* where negative autoregulation of *Ubx* (Hox) buffers gene expression to ensure appropriate haltere development ^101,102^. In the case of UNC-3, we show that both decreased and increased *unc-3* expression, or selective disruption of negative autoregulation, have consequences on MN molecular identity and animal locomotion. Because increased or decreased *EBF3* expression is associated with HADDS^7,8^, and syndromic variants disrupt *unc-3* negative autoregulation, our findings point to *EBF3* negative autoregulation as a possible pathogenic mechanism. Altogether, we propose that continuous and opposing regulatory inputs (HOX/PBX versus UNC-3) maintain *unc-3* expression within a critical range to secure postmitotic neuronal identity and function throughout life. A similar “balancing act” of negative autoregulation counteracted by positive input has been observed in vertebrate embryonic stem cells with the TF Oct4, where its precise expression levels are required to maintain stem cell self-renewal ^103,104^.

Maintenance of TF expression in post-mitotic cells is often accomplished via positive autoregulation ^20–25^. This mechanism applies to many TFs functioning in long-lived cell types - such as pancreatic acinar cells ^25^ and skeletal muscle ^23,24^. In neurons, positive autoregulation has been demonstrated for a handful of terminal selectors to date ^21,22,34,39,42,45–54^. Positive autoregulation can function to boost TF expression levels above a critical threshold required for cell identity acquisition ^22^. Positive autoregulation also functions to “lock-in” a cell fate after transient signals that induce TF expression have been lost, functioning as a bistable switch in a cell fate choice ^18^. Bistable switches emerging from positive autoregulation are reversible; molecular fluctuations in TF levels can cause sufficient loss of the positive autoregulatory input, leading to loss of a particular cell fate ^55,56^. As a result, additional mechanisms, such as epigenetic remodeling ^57^ and additional TF network motifs ^50,56^, are needed to stabilize cell fate. Because negative autoregulation can function to restrict gene expression variability ^30^, we surmise that it provides a mechanism for cell fate maintenance, ensuring robustness against environmental or developmental noise.

Altogether, we propose that regulation of dosage-sensitive TFs over time through a ‘balancing-act’ of activating and self-repressing inputs may constitute a general principle for cell fate maintenance in long-lived cell types. Because *EBF3* syndromic variants disrupt *unc-3* negative autoregulation, our findings are relevant for our understanding of HADDS and other disorders caused by mutations or variation in dosage-sensitive TFs.

## Supporting information

Suppl. Table 1. Structural predictions of missense mutations

Suppl. Table 2. Worm strains used in this study

Suppl. Table 3. Primers used in this study.

## ACKNOWLEDGMENTS

We thank Ilaria Rebay, Ellie Heckscher, Alex Ruthenberg, Oliver Hobert, Sumin Jang and members of the Kratsios lab (Ian Weigle, Filipe Marques, Nidhi Sharma) for comments on the manuscript. We also thank Harry Feng, Nimisha Krishnan, Isabel Layne, and Ruby Baer for contributions to this work. We are grateful for Oliver Hobert’s support, as the initial observations on negative autoregulation were made by P.K in the Hobert lab at Columbia University. We acknowledge invaluable community sources, including: the *Caenorhabditis* Genetics Center (CGC), which is funded by NIH Office of Research Infrastructure Programs (P40 OD010440), for providing strains; WormBase ^105^, which was used for experimental design and execution; and the Encyclopedia of DNA Elements (ENCODE) and ChIP-Atlas for the publicly available CEH-20 ChIP-seq and neuronal ATAC-seq data sets, respectively ^78,90,106,107^. We thank the members of the Research Computing Center (RCC) at the University of Chicago for providing their services. This work was supported by an NSF fellowship (2140001) to H.D., and two NIH grants (R01 NS118078, R01 NS116365) to P.K.

## AUTHOR CONTRIBUTIONS

H.D., Conceptualization, data curation, investigation, formal analysis, visualization, methodology, writing – original draft, review, and editing; A.K., Investigation, formal analysis; H.S., Investigation, formal analysis. A.E.X. Investigation, formal analysis, supervision; P.K. Conceptualization, supervision, investigation, funding acquisition, project administration, writing – original draft, review, and editing;

## DATA AVAILABILITY

ChIP-seq data for CEH-20 was downloaded from ENCODE (https://www.encodeproject.org/) (Waterston Lab, identifier: ENCFF249XSX). ChIP-seq data for UNC-3 is accessible on GEO (https://www.ncbi.nlm.nih.gov/geo/query/acc.cgi?acc=GDS) with accession number GSE143165. ChIP-seq data for EBF3 is accessible on GEO with accession number GSE90682. Neuronal ATAC-seq data is available on https://chip-atlas.org/peak_browser from the Ahringer Lab ^72^.

## MATERIALS AND METHODS

### C. elegans strains

Worms were grown at 15, 20, or 25°C on nematode growth media (NGM) plates seeded with bacteria (*E. coli* OP50) as food source. Table of all strains used in this study can be found in **Supplemental Table 2**. Genotyping primers are found in **Supplemental Table 3**.

### *C. elegans* staging, imaging, and fluorescence quantification

L2 worms of syb8054 strain were staged by animal length at L1/L2 molt (350 – 450 microns from nose to tail). L4 worms were staged by vulval morphology (Christmas tree stage). For UNC-3::GFP expression profile over time, worms were staged by the following methods: (L1) laid eggs were picked to a separate plate and worms were imaged as they hatched. (L2) birth of larvally born motor neurons. (L3) number of vulval precursor cells. (L4) vulval morphology at Christmas tree stage. (Young Adults) presence of 1-4 embryos. (Day 1 and 2 adults) L4 worms were picked to a separate plate and imaged after 24 and 48 hours, respectively.

Worms were mounted on a 4% agarose pad on a glass slide and anesthetized with 100 mM sodium azide (NaN_3_). Fluorescence images were captured with a Zeiss LSM 900 confocal or a Zeiss Axio Imager.Z2 epifluorescent microscope. Z-stacks were acquired in the ZEN software.

Representative images are maximum intensity projections, and all image reconstruction and analyses were performed in Fiji ^108^. Background subtraction was done with a rolling ball radius of 50 pixels using a sliding paraboloid. Pixel intensity was quantified in the Z slice of greatest in-focus intensity for each neuron.

### Motor neuron identification

Motor neurons were identified based on a combination of: (i) invariant cell body positioning along the ventral nerve cord, (ii) localization relative to fluorescent GABA-ergic motor neuron markers, and (iii) motor neuron birth order.

### Protein domain analysis

Intrinsically disordered regions were identified using Uniprot and four separate programs: Predictors of Natural Disordered Regions (PONDR), Intrinsically Unstructured Proteins Predictor 2 ANCHOR (IUPred2A) ^109^, putative Function- and Linker based Disorder Prediction using deep Neural Network (flDPnn) ^110^, and NetSurfP-3.0 ^111^. An intrinsically disordered region was defined as a region where at least three of the five resources called residues disordered. Protein domains were aligned using EMBOSS Needle Pairwise Sequence Alignment ^112^.

### Identification of COE binding sites

UNC-3 binding sites (COE motifs) were predicted using the Find Individual Motif Occurrences (FIMO) from MEME suite ^113^, with a *p-*value threshold of *p* < .001. The position weight matrix (PWM) for the UNC-3 binding site was previously derived as described here ^61^.

### Generation of transgenic animals

The fosmid-based *unc-3* reporter was generated through recombineering using fosmid clone WRM0622bH08, coordinates chrX:14,760,199-14,791,825, and was injected as a complex array with the co-injection marker myo-2::GFP (3ng/uL) ^114^.

Extrachromosomal constructs for *unc-3* cDNA expression and for *cis*-regulatory mutant analysis were generated by Gibson Assembly (NEB catalog #E5510S). For *unc-3* cDNA constructs, linear purified PCR products of the *unc-3*_cDNA construct (15 ng/uL) were injected into *kasIs13 ; unc-3(e151)* worms with a co-injection marker of Pttx-3::mCherry (50 ng/uL). For *cis-*regulatory mutant analysis, COE site mutation was generated via PCR mutagenesis. Linear purified PCR products (30 ng/uL) were injected into N2 worms with a co-injection marker of Pttx-3::GFP (30 ng/uL). For each extrachromosomal construct, 2-3 independent lines were generated from separate F1s arising from the injected parent that then gave rise to progeny stably inheriting the array.

Promoter sequences and primers can be found in **Supplemental Table 3.**

### Genome engineering using CRISPR-Cas9

CRISPR/Cas9 genome editing was conducted following Ghanta & Mellow (2020) protocol ^115^. Briefly, injection mix was prepared with Cas9 (0.5 µL of 10 µg/µL stock), tracrRNA (5 µL of 0.4 µL/µg stock), and crRNA (2.8 µL of 0.4 µg/µL stock) and incubated at 37 degrees Celsius for 15 minutes before addition of ssODN (2.2 µL of 1 µg/µL), co-injection marker (rol-6, 1.6 µL of 500 ng/µL stock), and nuclease free water (up to 20 µL).

crRNA and ssODN sequences used can be found in **Supplemental Table 3**.

Additional CRISPR-edited strains were generated by SUNYbiotech (see **Supplemental Table 2**).

### Temporally controlled protein degradation

Auxin-induced protein degradation was conducted as published ^70^. Briefly, auxin-tagged alleles (ceh-20(syb6724[ceh-20::3XFLAG::mNG::AID]) III and unc-3(ot837[unc-3::mNG::2xFLAG::AID])) were crossed with a ubiquitously expressed TIR-1 (ieSi57 [eft-3p::TIR1::mRuby::unc-54 3’UTR + Cbr-unc-119(+)] II) and exposed to natural (indole-3-acetic acid [IAA]) or synthetic (1-napthalenceacetic acid, potassium salt [K-NAA]) auxins. IAA (dissolved in EtOH) or K-NAA (dissolved in diH_2_O) was added to NGM plates for a final concentration of 4 mM. Plates were seeded with OP50 and shielded from light.

### Thrashing assays

To assess animal thrashing, worms were placed in a 40 uL drop of M9 on an NGM plate with no food at room temperature. After 1 minute of acclimatization, worms were filmed under a stereo microscope for 30 seconds. The video was slowed to half speed to count the number of body bends (turning of the head and tail to the same side of the body) within 30 seconds.

### Behavioral tracking

Worms were maintained as mixed stage populations with *E. coli* OP50 at 20°C until bleaching synchronization. 10 – 15 day 1 adult worms were then transferred to 96 square well plates (Whatman UNIPLATE 96-square well plate, WHAT-77011651) with 200 uL of NGM agar seeded with 10 µL of PFA treated *E. coli* OP50. Worms on the plates were acclimatized for 30 minutes prior to imaging at 20 degrees in the dark. Worms were tracked and the videos were processed as previously described ^116^. Three videos were recorded sequentially, a 5 min pre-stimulus recording, followed by a 6 min blue bight stimulation recording containing three 10 s blue light pulses delivered at 60 s, 160 s, and 260 s, and finally a 5 min post-stimulus recording. Recordings were controlled using a script written with LoopBio’s Motif API (https://github.com/loopbio/python-motifapi; nzjrs, 2024). Four biological replicates were performed for each worm strain, with each biological replicate corresponding to one 96 square well imaging plate.

### Behavioral feature extraction and analysis

Videos were then segmented and animals tracked using Tierpsy Tracker ^117^. Following skeleton segmentation, a previously described convolutional neural network classifier was applied to exclude non worm objects prior to feature extraction (Barlow et al., 2022). Skeletons that did not meet the morphological criteria were excluded from analysis, 700–1300 µm in length and 20–200 µm in width. Following tracking, a previously defined set of 3,076 behavioural features was extracted from each well from each of the three recordings (pre-stimulus, blue-light stimulation, and post-stimulus) ^86^. Feature values were averaged across all tracks to generate a single feature vector for each well. Pairwise comparisons between each treatment class and control were performed using two-sided t-test. Multiple testing correction was applied per class across all features using fdr with alpha = 0.05. Hierarchical clustering was performed using ward linkage and euclidean distance on standardized feature matrices. Isolation Forest analysis was conducted using N/A bootstrap samples per iteration, contamination rate 0.2, 20 iterations, and a consensus threshold of 80%.

### *In silico* protein-protein interaction screen and missense predictions

A local implementation of Alphafold2 ^118,119^, LocalColabFold ^82^ (https://github.com/YoshitakaMo/localcolabfold), was used to predict protein-protein interactions. Multiple sequence alignments were generated using MMseqs2 ^120^. 5 recycling steps were used. Likely candidates were identified as those where the ipTM score was above 0.3 for at least 3 of the 5 models, as previously observed ^63,85^. Top ranked models were visualized using PyMOL (https://pymol.org/).

Effects of missense mutations were determined by structure modelling with Alphafold3 with default setting (https://alphafoldserver.com/) ^121^. UNC-3 was modeled as a dimer with 130bp of the *unc-17* promoter sequence containing a palindromic COE binding site (TCCCCGGGGA). Accurate prediction of interaction between the UNC-3 dimer and the COE binding site was verified using PDBsum (https://www.ebi.ac.uk/thornton-srv/databases/pdbsum/) and compared with interactions identified in the EBF1 crystal structure ^73^)\. Models and polar contacts were assessed using PyMol (https://pymol.org/).

### Statistical analysis and reproducibility

Statistical analyses were conducted in RStudio and graphs were generated with ggplot. Data were plotted with jitter. Boxplots depict 25^th^ and 75^th^ percentiles (hinges) and median (center bar), with whiskers extending to minimum and maximum values.

