## Supplemental Figures 1- 6 for "Continuous negative autoregulation fine-tunes dosage-sensitive transcription factor expression to maintain post-mitotic neuron identity"

### Supplemental Figure 1

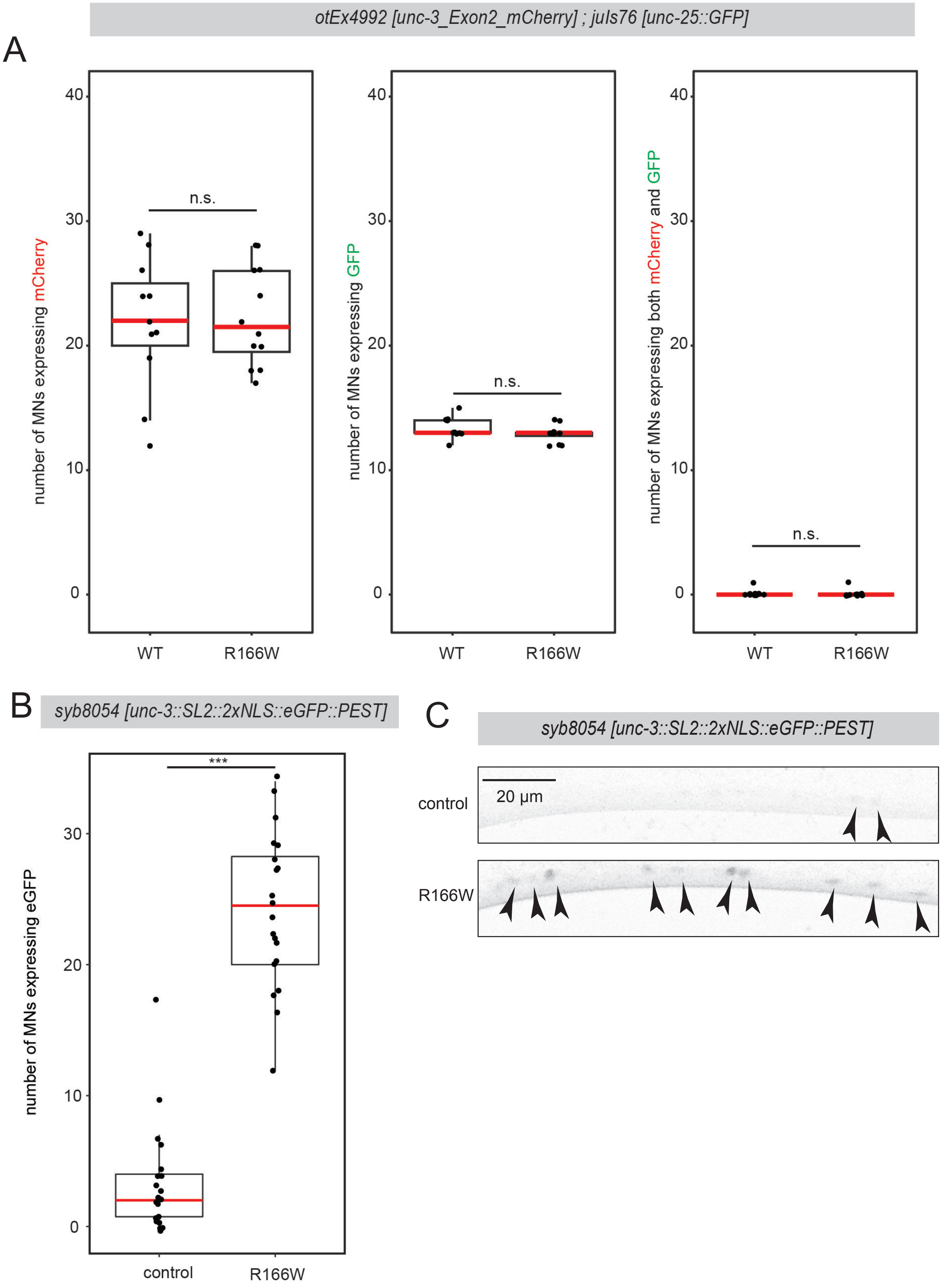


**Supplemental Figure 1:** *unc-3* negatively autoregulation occurs in cholinergic motor neurons, related to Figure 1.
**A:** Quantification of number of motor neurons (MNs) expressing fosmid-based transcriptional *unc-3* reporter (left), transgenic GABA marker *unc-25* reporter (middle), or both (right) in N2 or *unc-3(R166W)* backgrounds at L4. Dots represent individual worms. Red bar represents median. Welch's T-test. (N2 n = 11 worms, *unc-3(R166W)* n = 12) **B:** Quantification of number of MNs expressing *unc-3* endogenous transcriptional reporter at L4 in control or *unc-3(R166W)* backgrounds. Dots represent individual worms. Red bar represents median. Welch's T-test. (control n = 11 worms, *unc-3(R166W)* n = 12) **C:** Representative images of *unc-3* endogenous transcriptional reporter at L4 in control or *unc-3(R166W)* backgrounds. Arrowheads: motor neurons. Scale bar: 20 µm. p-values: * < .05, ** <.01, *** < .001, n.s. = no significance.

### Supplemental Figure 2

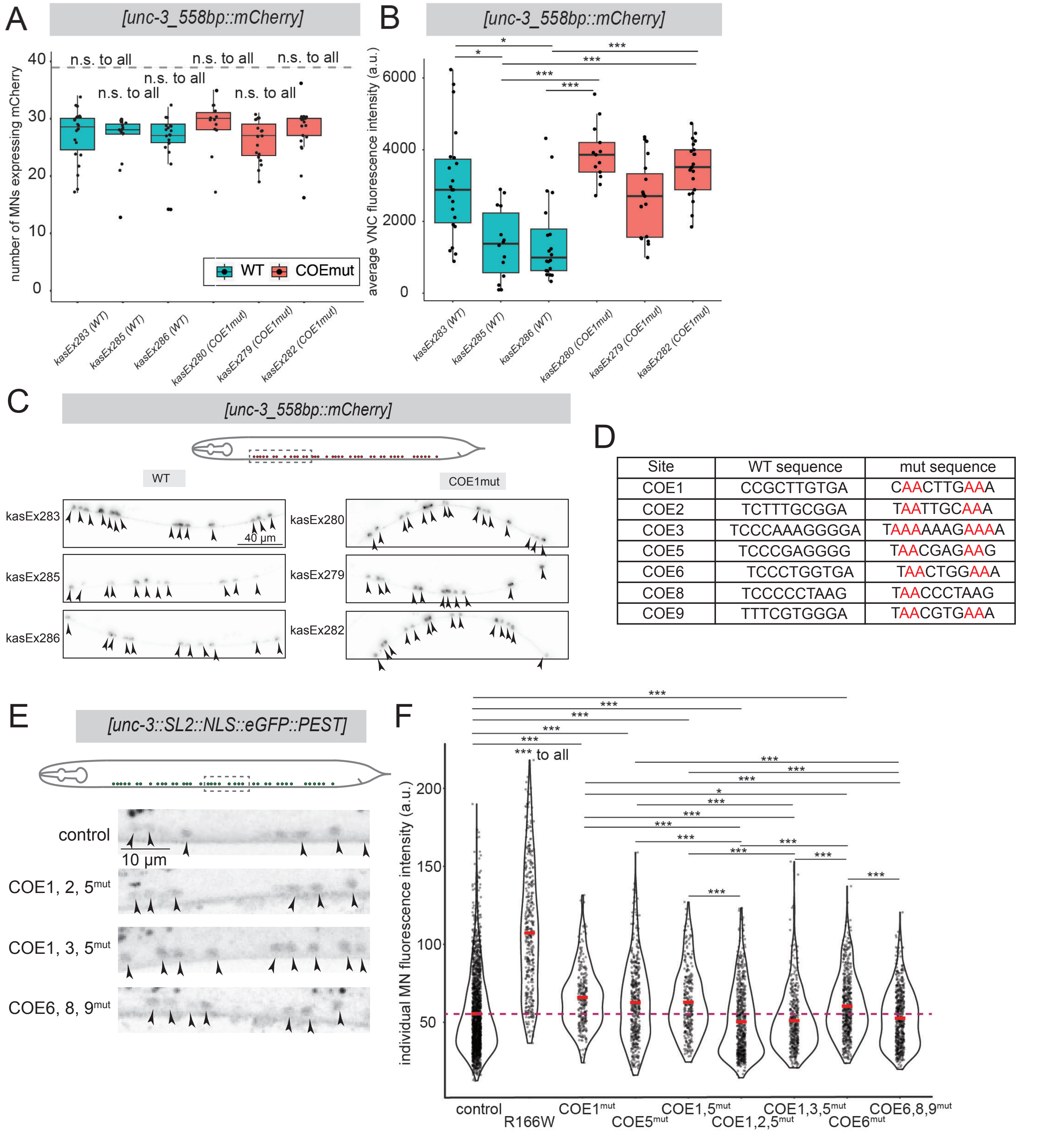


| Site | WT sequence | mut sequence |
|---|---|---|
| COE1 | CCGCTTGTGA | CAACTTGAAA |
| COE2 | TCTTTGCGGA | TAATTGCAAA |
| COE3 | TCCCAAAGGGGA | TAAAAAAGAAAA |
| COE5 | TCCCGAGGGG | TAACGAGAAG |
| COE6 | TCCCTGGTGA | TAACTGGAAA |
| COE8 | TCCCCCTAAG | TAACCCTAAG |
| COE9 | TTTCGTGGGA | TAACGTGAAA |

**Supplemental Figure 2:** *unc-3* negative autoregulation is direct through certain combinations of COE sites, related to Figure 2.
**A:** Quantification of number of motor neurons (MNs) expressing transgenic transcriptional *unc-3* reporter with wild-type COE1 (blue) or COE1$^{mut}$ (orange) sequences at L4. Middle black bar represents median. Dashed grey line represents the 39 total cholinergic MNs in the ventral nerve cord. Kruskal-Wallis multiple comparisons test with Bonferroni correction. (*kasEx283* n = 22 worms, *kasEx285* n = 14, *kasEx286* = 20, *kasEx280* = 13, *kasEx279* = 19, *kasEx282* = 21). **B:** Quantification of average MN fluorescence intensity of transgenic transcriptional *unc-3* reporter with wild-type COE1 (blue) or COE1$^{mut}$ (orange) sequences at L4, subdivided by independent lines. Middle black bar represents median. Non-significant comparisons not shown. (*kasEx283* n = 22 worms, *kasEx285* n = 14, *kasEx286* n = 20, *kasEx280* n = 13, *kasEx279* n = 19, *kasEx282* n = 21). **C:** Representative images of transcriptional *unc-3* reporter with wild-type COE1 or mutant COE1 sequences at L4, subdivided by independent lines. Top: Schematic of nerve cord section displayed in images. Arrowheads: MNs. Scale bar: 40 µm **D:** Table of COE site sequences and their respective mutations in this study. **E:** Representative images of *unc-3* endogenous transcriptional reporter with control, COE1,2,5$^{mut}$, COE1,3,5$^{mut}$, or COE6,8,9$^{mut}$ backgrounds at L2. Top: Schematic of nerve cord section displayed in images. Arrowheads: MNs. Scale bar: 10 µm **F:** Quantifications of individual MN fluorescence intensities of *unc-3* endogenous transcriptional reporter with control, R166W, COE1$^{mut}$, COE5$^{mut}$, COE1,5$^{mut}$, COE1,2,5$^{mut}$, COE1,3,5$^{mut}$, COE6$^{mut}$, or COE6,8,9$^{mut}$ backgrounds at L2. Dots represent individual MNs. Red bar represents mean. Dunn Kruskal-Wallis multiple comparisons test with Bonferroni correction. (control n = 74 worms, unc-3(R166W) n = 12, COE1$^{mut}$ n = 9, COE5$^{mut}$ n = 17, COE1 and 5$^{mut}$ n = 14, COE1,2,5$^{mut}$ n = 25, COE1,3,5$^{mut}$ n = 15, COE6$^{mut}$ n = 24, COE6,8,9$^{mut}$ n = 25). Purple dashed line represents the mean of the control. Non-significant values not shown. p-values: * < .05, ** <.01, *** < .001, n.s. = no significance.

### Supplemental Figure 3

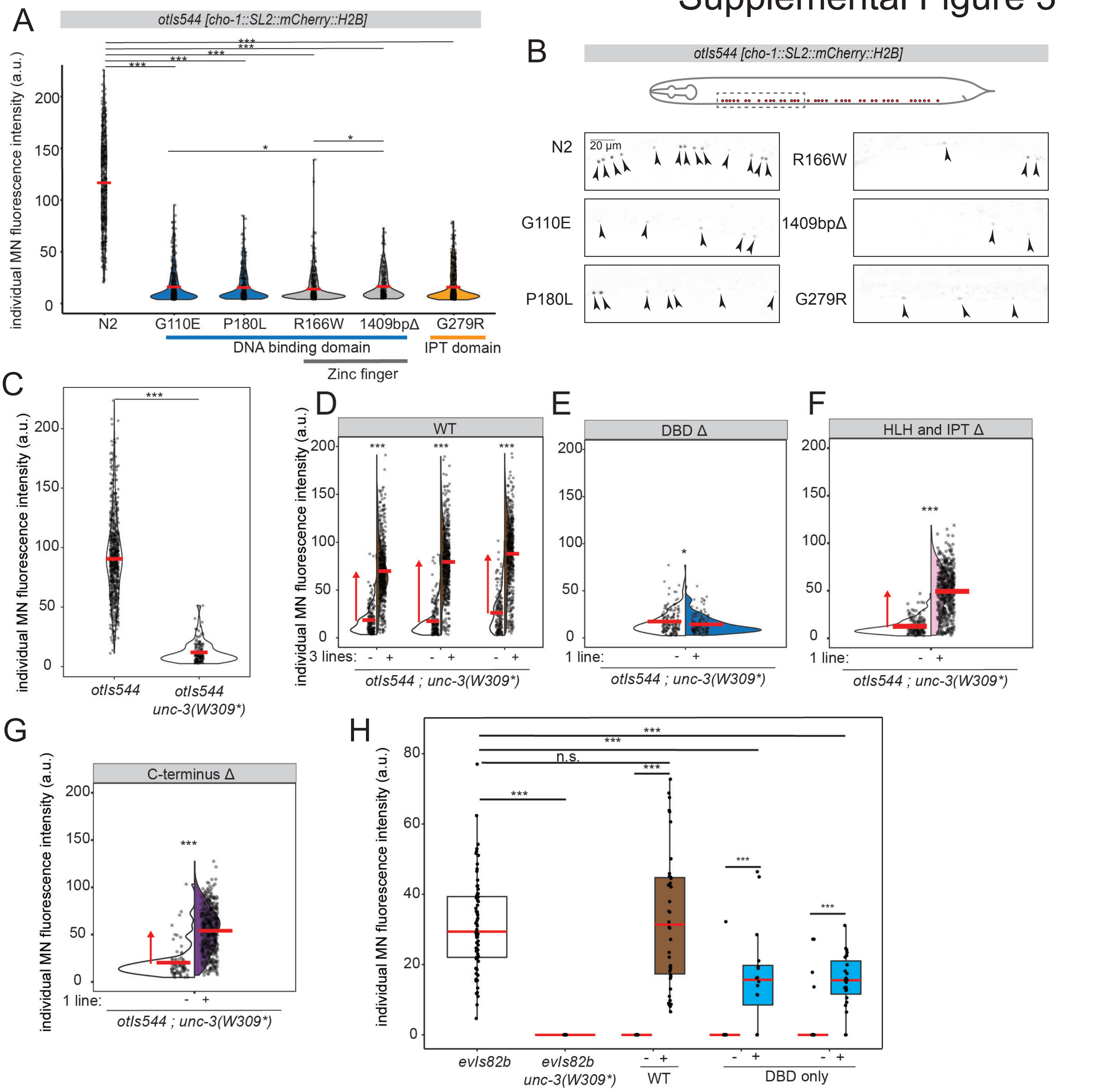


**Supplemental Figure 3:** The DNA-binding and IPT domains are required for target gene activation, related to Figure 5.

**A:** Quantification of individual motor neuron (MN) fluorescence intensity of fosmid-based transcriptional *cho-1* reporter at L4 in N2, *unc-3(G110E), unc-3(P180L), unc-3(R166W), unc-3(1409bpΔ)*, and *unc-3(G279R)* backgrounds. Dots represent individual MN fluorescence intensities. Red bar represents mean. Domains are colored as: blue – DNA binding domain, grey – Zinc finger, orange – Immunoglobulin/Plexin/Transcription factor (IPT) domain. Dunn Kruskal-Wallis multiple comparisons test with Bonferroni correction. (N2 n =15 worms, *unc-3(G110E)* n = 17, *unc-3 (P180L)* n = 17, *unc-3(R166W)* n = 17, *unc-3(1409bpΔ)* n = 12, *unc-3(G279R)* n = 19). **B:** Representative images of fosmid-based transcriptional *cho-1* reporter at L4 in N2, *unc-3(G110E), unc-3(P180L), unc-3(R166W), unc-3(1409bpΔ)*, and *unc-3(G279R)* backgrounds. Top: Schematic of nerve cord section displayed in images. Arrowheads: MNs. Scale bar: 20 μm **C:** Quantification of individual MN fluorescence intensity of fosmid-based transcriptional *cho-1* reporter at L4 in N2 and *unc-3(W309*)* backgrounds. Dots represent individual MN fluorescence intensities. Red bar represents mean. Welch's T-test. (control n = 21 worms, *unc-3(W309*)* n = 19) **D-G:** Quantification of individual MN fluorescence intensity of fosmid-based transcriptional *cho-1* reporter at L4 in *unc-3(W309*)* background either with (right, colored) or without (left, uncolored) the respective rescue construct (top). Dots represent individual MN fluorescence intensities. Red bar represents mean. Red arrow highlights mean change in construct-carrying vs non-construct-carrying siblings. Construct consists of *Punc-3_558bp* driving unc-3 cDNA (D) WT (*kasEx357*+ n = 17, *kasEx357*- n = 15, *kasEx358*+ n = 18, *kasEx358*- n = 18, *kasEx356*+ n = 17, *kasEx356*- n = 20), (E) DBD deletion (*kasEx364*+ n = 19 worms, *kasEx364*- n = 20) (F) combined Helix Loop Helix (HLH) and IPT domain deletion (*kasEx370*+ n = 20 worms, *kasEx370*- n = 19), (G) C-terminus deletion (*kasEx375*+ n = 17, *kasEx375*- n = 15). Welch's T-test. **H:** Quantification of individual MN fluorescence intensity of *unc-129* reporter at L4 in N2 and *unc-3(W309*)* backgrounds, and in *unc-3(W309*)* background either with (right, colored) or without (left, uncolored) the respective rescue construct. Brown: WT (*kasEx388*+ n 14 = , *kasEx388*- n = 14); Bright blue: DBD only (*kasEx390*+ n = 13, *kasEx390*- n = 18, *kasEx389*+ n = 12, *kasEx389*- n = 19). Dots represent individual MN fluorescence intensities. Red bar represents mean. Welch's T-test. (control n = 15 worms, *unc-3(W309*)* = 14). p-values: * < .05, ** <.01, *** < .001, n.s. = no significance.

### Supplemental Figure 4

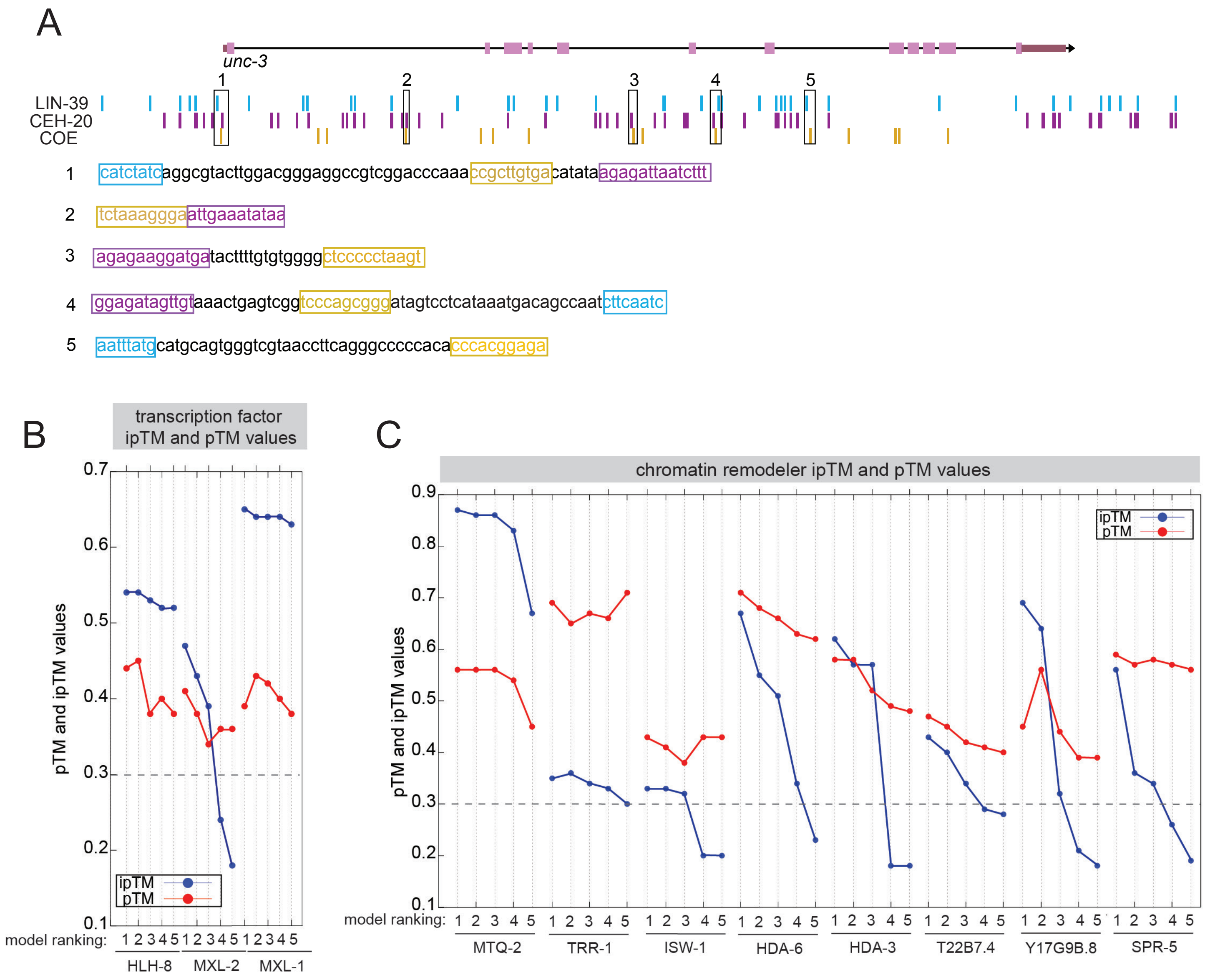


D

| | *C. elegans* gene | human orthologue |
|---|---|---|
| Transcription factors | HLH-8 | TWIST |
| | MXL-1 | MAX |
| | MXL-2 | MLX |
| Chromatin associated factors | HDA-3 | HDAC1-2 |
| | HDA-6 | HDAC6 |
| | ISW-1 | SMARCA1 |
| | MTQ-2 | HEMK2 |
| | T22B7.4 | SGF29 |
| | TRR-1 | TRRAP |
| | SPR-5 | KDM1A |
| | Y17G9B.8 | SGF29 |

**Supplemental Figure 4:** An *in silico* screen identifies chromatin factors and transcription factors as putative UNC-3 cofactors, related to Figure 5.
**A:** Schematic of *unc-3* locus and bioinformatically identified binding sites for LIN-39 (blue), CEH-20 (purple), and UNC-3 (yellow). Binding site p-value < .001. 5 insets: zoom in to COE sites in close proximity to activator binding sites to show their spacing.
**B-C:** ColabFold pTM (red) and ipTM (blue) values for transcription factor **(B)** and chromatin remodeler **(C)** dimerization with UNC-3, 5 models each. pTM: predicted template modelling. ipTM: interface predicted template modelling. Gray dashed line: 0.3 ipTM cutoff for warranting further investigation.
**D:** Table of 11 identified candidates and their respective orthologues, sorted by transcription factors and chromatin associated factors.

### Supplemental Figure 5

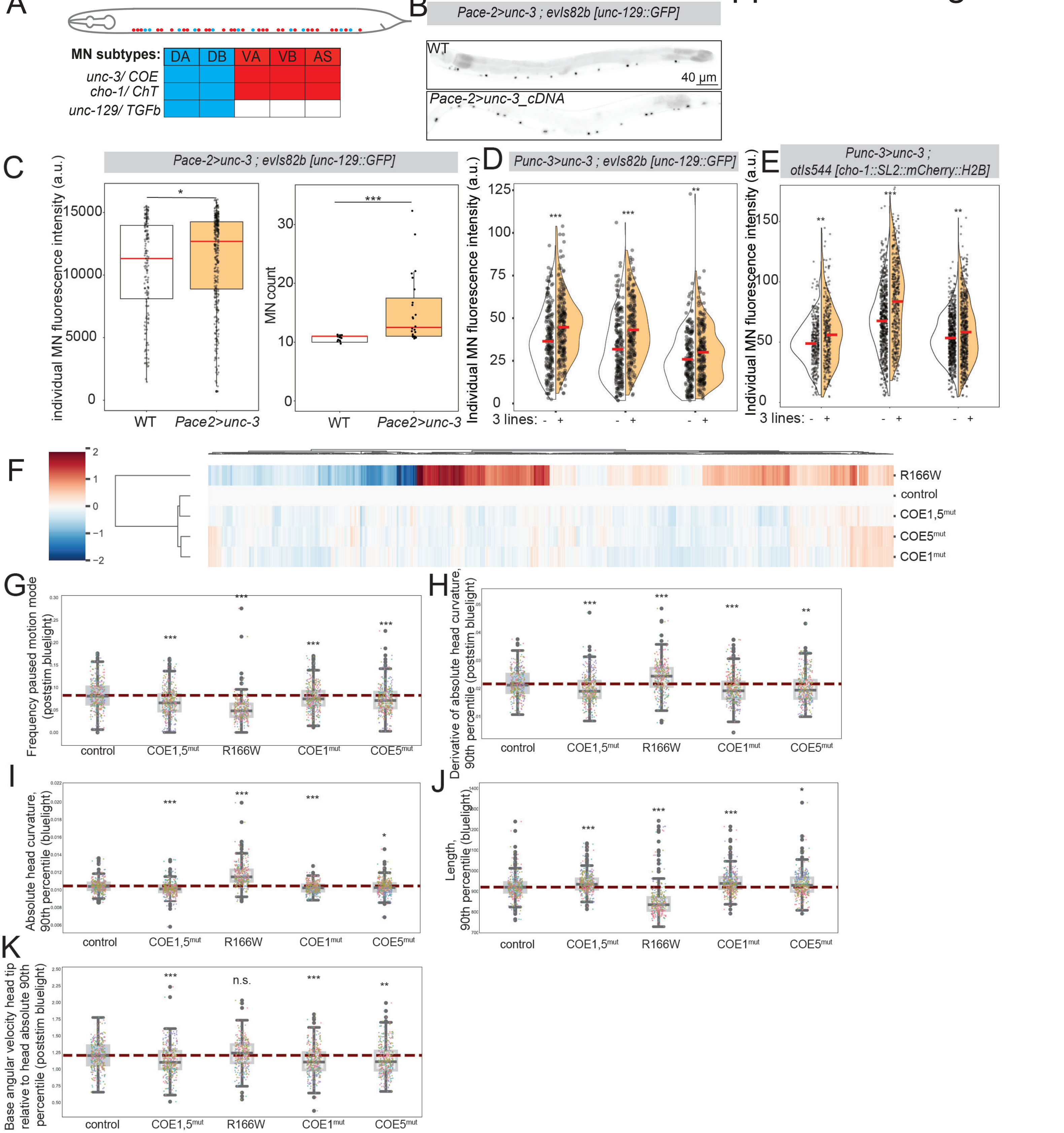


**Supplemental Figure 5:** UNC-3 dosage-dependent effects on motor neuron identity genes and behavior, related to Figure 6.
**A:** Schematic of UNC-3+ motor neuron (MN) subtypes and terminal identity gene expression. **B:** Representative images of *unc-129::GFP* reporter in control (top) or *ace-2p>unc-3_cDNA* (bottom) **C:** Quantification of individual MN fluorescence intensity (left) and number of GFP+ MNs (right) of *unc-129::GFP* reporter in control or *ace-2p>unc-3_cDNA*. Dots represent individual MN fluorescence intensities (left) and individual worms (right). Red bar represents mean. Welch's T-test. (control n = 14 worms , *ace-2p>unc-3* n = 14) **D:** Quantification of individual MN fluorescence intensity of transgenic *unc-129::GFP* reporter at L4 either with (right, colored) or without (left, uncolored) *Punc-3_558bp>unc-3_cDNA*. Dots represent individual MN fluorescence intensities. Red bar represents mean. Welch's T-test. (*kasEx356+* n = 20 worms, *kasEx356-* n = 19, *kasEx358+* n = 19, *kasEx358-* n = 20, *kasEx357+* n = 20, *kasEx357-* n = 20) **E:** Quantification of individual MN fluorescence intensity of transcriptional *cho-1* reporter at L4 either with (right, colored) or without (left, uncolored) *Punc-3_558bp>unc-3_cDNA*. Dots represent individual MN fluorescence intensities. Red bar represents mean. Welch's T-test. (*kasEx356+* n = 9 worms, *kasEx356-* n = 9, *kasEx358+* n = 12, *kasEx358-* n = 19, *kasEx357+* n = 18, *kasEx357-* = 20) **F:** Hierarchically clustered heatmap of 256 features of automated worm tracking. **G-K:** Quantification of **(G)** frequency of paused motion post blue light stimulation, **(H)** derivative of the 90th percentile of head curvature post blue light stimulation, **(I)** 90th percentile of the curvature of the head during blue light stimulation, **(J)** 90th percentile of length, **(K)** 90th percentile of angular velocity of the head tip post blue light stimulation. Welch's T-test. Outliers shown in open circles. Solid red bar represents median. Dashed red bar represents mean of the control. p-values: * < .05, ** <.01, *** < .001, n.s. = no significance.

### Supplemental Figure 6

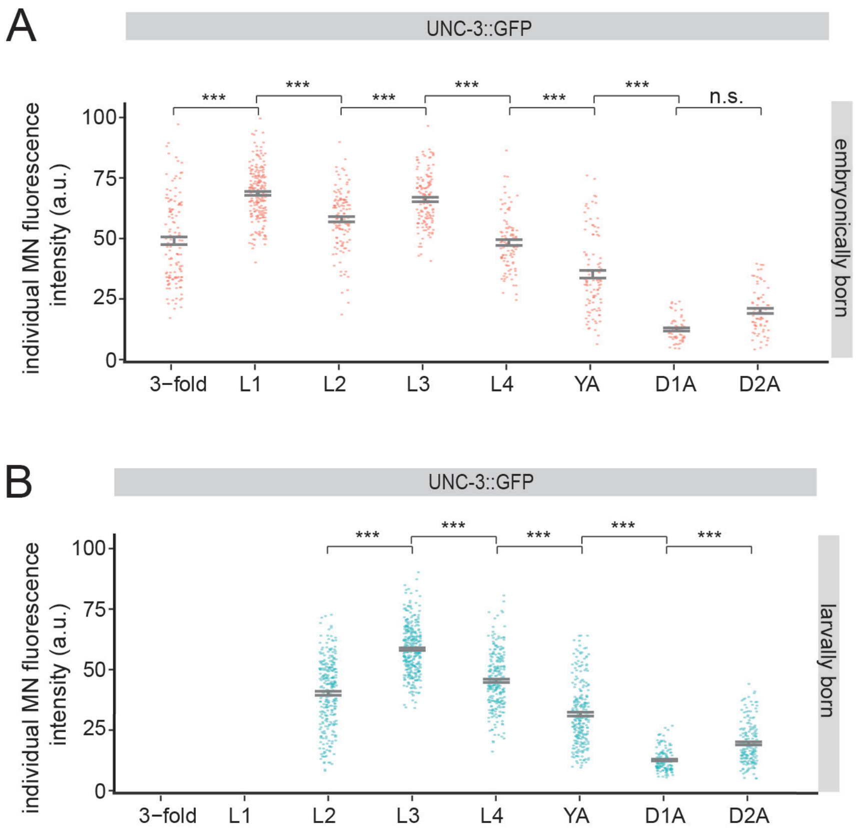


**Supplemental Figure 6:** UNC-3 expression is continuous over time, related to Figure 7.
**A-B:** Quantification of individual motor neuron (MN) fluorescence intensity of UNC-3 endogenous translational reporter across time, for **(A)** embryonically born and **(B)** larvally bornMNs. Dots represent individual MN fluorescence intensities. Standard error bars shown in grey. Tukey honest significance test. (3-fold n = 12 worms, L1 n = 18, L2 n = 14, L3 n = 15, L4 n = 13, YA n = 12, D1A n = 8, D2A n = 9).
p-values: * < .05, ** <.01, *** < .001, n.s. = no significance.
